# Therapeutic immune cell engineering with an mRNA : AAV-*Sleeping Beauty* composite system

**DOI:** 10.1101/2023.03.14.532651

**Authors:** Lupeng Ye, Stanley Z. Lam, Luojia Yang, Kazushi Suzuki, Yongji Zou, Qianqian Lin, Yueqi Zhang, Paul Clark, Lei Peng, Sidi Chen

## Abstract

Adoptive cell therapy has shown clinical success in patients with hematological malignancies. Immune cell engineering is critical for production, research, and development of cell therapy; however, current approaches for generation of therapeutic immune cells face various limitations. Here, we establish a composite gene delivery system for the highly efficient engineering of therapeutic immune cells. This system, termed MAJESTIC (**<u>m</u>**RNA **A**AV-Sleeping-Beauty **<u>J</u>**oint **<u>E</u>**ngineering of **<u>S</u>**table **<u>T</u>**herapeutic **<u>I</u>**mmune **<u>C</u>**ells), combines the merits of mRNA, AAV vector, and transposon into one composite system. In MAJESTIC, the transient mRNA component encodes a transposase that mediates permanent genomic integration of the *Sleeping Beauty* (SB) transposon, which carries the gene-of-interest and is embedded within the AAV vector. This system can transduce diverse immune cell types with low cellular toxicity and achieve highly efficient and stable therapeutic cargo delivery. Compared with conventional gene delivery systems, such as lentiviral vector, DNA transposon plasmid, or minicircle electroporation, MAJESTIC shows higher cell viability, chimeric antigen receptor (CAR) transgene expression, therapeutic cell yield, as well as prolonged transgene expression. CAR-T cells generated by MAJESTIC are functional and have strong anti-tumor activity *in vivo*. This system also demonstrates versatility for engineering different cell therapy constructs such as canonical CAR, bi-specific CAR, kill switch CAR, and synthetic TCR; and for CAR delivery into various immune cells, including T cells, natural killer cells, myeloid cells, and induced pluripotent stem cells.

## Introduction

Cellular immunotherapy involves the administration of “living drugs”: genetically modified immune cells that can proliferate, adapt to their environment, engage surrounding cells, and elicit dynamic responses that directly or indirectly target tumor cells for destruction ^1^. Adoptive cell transfer (ACT) is one type of cellular immunotherapy which involves the transfer of cells that directly target tumor cells in the patient ^2^. One notable ACT approach is chimeric antigen receptor (CAR) T cell therapy, in which T cells are engineered to express a synthetic membrane receptor specific for a tumor antigen. CAR-T therapy has had a remarkable effect in patients with certain hematological malignancies ^3, 4^, with five CAR-T products currently approved by the U.S. Food and Drug Administration (FDA) for the treatment of multiple myeloma and B-cell malignancies ^5^. To tackle the myriad challenges that different tumors and tumor environments present, multiple cell-based therapies besides CAR-Ts have been generated, such as tumor infiltrating lymphocytes (TILs) ^6^, T cell receptor engineered T (TCR-T) cells ^7, 8^, CAR-NK cells ^9–15^, CAR macrophages (CAR-Ms) ^16, 17^, and human induced pluripotent stem cell (iPSC)-derived therapeutic immune cells ^15 17, 18^. Additional components have been added to CAR constructs to enhance therapeutic efficacy or safety, e.g., kill switch-CARs that can be depleted upon administration of a drug in the case of deleterious CAR toxicity ^19^ or tandem CARs that can target two different antigens ^20^. A vital part of the implementation of such cellular immunotherapies is the therapeutic cell generation process: whether and how stably engineered therapeutic immune cells can be efficiently generated has a critical impact on cell therapy ^21^.

The majority of current CAR-Ts used in clinical trials were generated using γ-retrovirus or lentivirus for gene transfer. However, several limitations of this family of viral vectors exist. For example, transgene expression may be silenced or dampened^22^. A key limitation of γ-retroviruses specifically is their preference of integration into promoters of active genes^23^, including proto-oncogenes as observed in a clinical trial ^24–26^. Genomic studies indeed have found that the insertion profile of γ-retroviruses is associated with a higher likelihood of oncogenic transformation ^21^. HIV-derived lentiviral vectors are considered safer than γ-retroviral vectors based on current studies ^27^, but their pathogenic origin and biosafety level 2 or 2+ (BSL2/BSL2+) classification still warrant caution.

Adeno-associated virus (AAV) is a single stranded DNA virus. AAV is commonly used in gene therapy and is considered a safer vector^28^ (consistent with its BSL1 classification). However, AAV alone is not an ideal CAR carrier - it is non-integrating and CAR expression is thus rapidly diluted during T cell expansion. While AAV can serve as a template for efficient generation of stably integrated CAR-T cells^29, 30^, such strategies require genome editing via tools such as CRISPR/Cas. Recent studies^31–33^ have shown chromosomal rearrangement or loss following CRISPR/Cas9-mediated genome editing.

Non-viral systems have been proposed to eliminate the need for viral vectors. However, all current non-viral cell therapy approaches also have their limitations. mRNA electroporation is another CAR-generation strategy, but the duration of CAR expression is extremely short due to the instability of mRNA ^21^. Transposon systems such as *PiggyBac*^34, 35^ and *Sleeping Beauty* (*SB*) ^36^ can be used to directly integrate DNA sequences into the genome - this is an advantage over lentiviruses and retroviruses, which rely on the relatively error-prone reverse transcriptase enzyme to convert RNA into DNA^37^. Transposons are also attractive for their non-pathogenic origin ^21^. However, current transposon/transposase approaches for therapeutic cell generation rely predominantly on DNA electroporation or transfection of plasmids or mini-circles (MCs), for which efficiency of delivery varies between different studies, and cellular toxicity is usually high ^36, 38–41^. Additionally, repeated transposon mobilization by transposase may lead to continuous risks of insertional mutagenesis or local chromosomal rearrangements^42^.

As it stands, all current existing cell therapy engineering approaches, both viral and non-viral, have important limitations. Our goal is to develop a non-CRISPR gene delivery system that is capable of generating stably integrated therapeutic immune cells that 1) achieves efficient genomic integration without reliance on retroviruses or lentiviruses, 2) theoretically limits the extent of transposon mobilization, 3) achieves higher viability compared to current methods, and 4) independent of gene editing. Here we developed a prototype of such a platform, **<u>MAJESTIC</u>** (**<u>m</u>**RNA **A**AV-Sleeping-Beauty **<u>J</u>**oint **<u>E</u>**ngineering of **<u>S</u>**table **<u>T</u>**herapeutic **<u>I</u>**mmune **<u>C</u>**ells), which combines the merits of mRNA, AAV vector, and transposon into one composite system. In MAJESTIC, the mRNA component encodes a transposase that mediates a pulse of genomic integration of the *Sleeping Beauty (SB)* transposon, which carries genes-of-interest and is embedded inside the AAV vector. Plasmid DNA is more stable than mRNA and each copy may produce multiple copies of mRNA^43^; thus, supplying the transposase as mRNA may limit remobilization and subsequent chromosomal rearrangement in principle.

We tested this new system in standard laboratory settings, alone and side-by-side with conventional gene delivery systems, such as lentiviral vector or DNA transposons including plasmid DNA and MC DNA electroporation, measuring transduction efficiency, cell viability, therapeutic cell yield, and stability of transgene expression. We also demonstrated the system’s versatility for engineering different cell therapy constructs such as canonical CAR, bi-specific CAR, kill switch CAR and synthetic T cell receptor (TCR), and for therapeutic cell engineering of various immune cell types including T cells, natural killer (NK) cells, pluripotent stem cells (iPSCs), and cells in the myeloid lineage.

## Results

### Establishment of the MAJESTIC system and high efficiency generation of CAR-T cells

The MAJESTIC system has two core components: 1) the AAV-SB vector carrying desired cell therapy transgenes (AAV-SB-CTx) and 2) mRNA encoding the SB transposase (mRNA-Transposase) (**Fig. 1a**). To generate the AAV-SB-CTx plasmid, we first established the AAV-SB plasmid by cloning the SB transposon, which is flanked by inverted repeats/direct repeats (IR/DR), in between the inverted terminal repeats (ITRs) of the AAV plasmid backbone ^44^. Into this chimeric AAV-SB backbone, single scFv CAR, tandem scFv CAR, TCR, suicide-gene CAR (CAR.iCasp9) constructs were cloned **(Fig. 1a; Fig. S1a)**. The hyperactive *SB* transposase SB100x ^45^ was delivered to cells via mRNA electroporation to facilitate genomic integration of the *SB* transposon construct, which was delivered via AAV transduction **(Fig. 1a)**. Importantly, this two-component design achieves integration of the *SB* transposon while limiting remobilization compared to that for plasmid transfection because the mRNA encoding the transposase is in principle less stable than plasmid DNA transposase.

**Figure 1.**
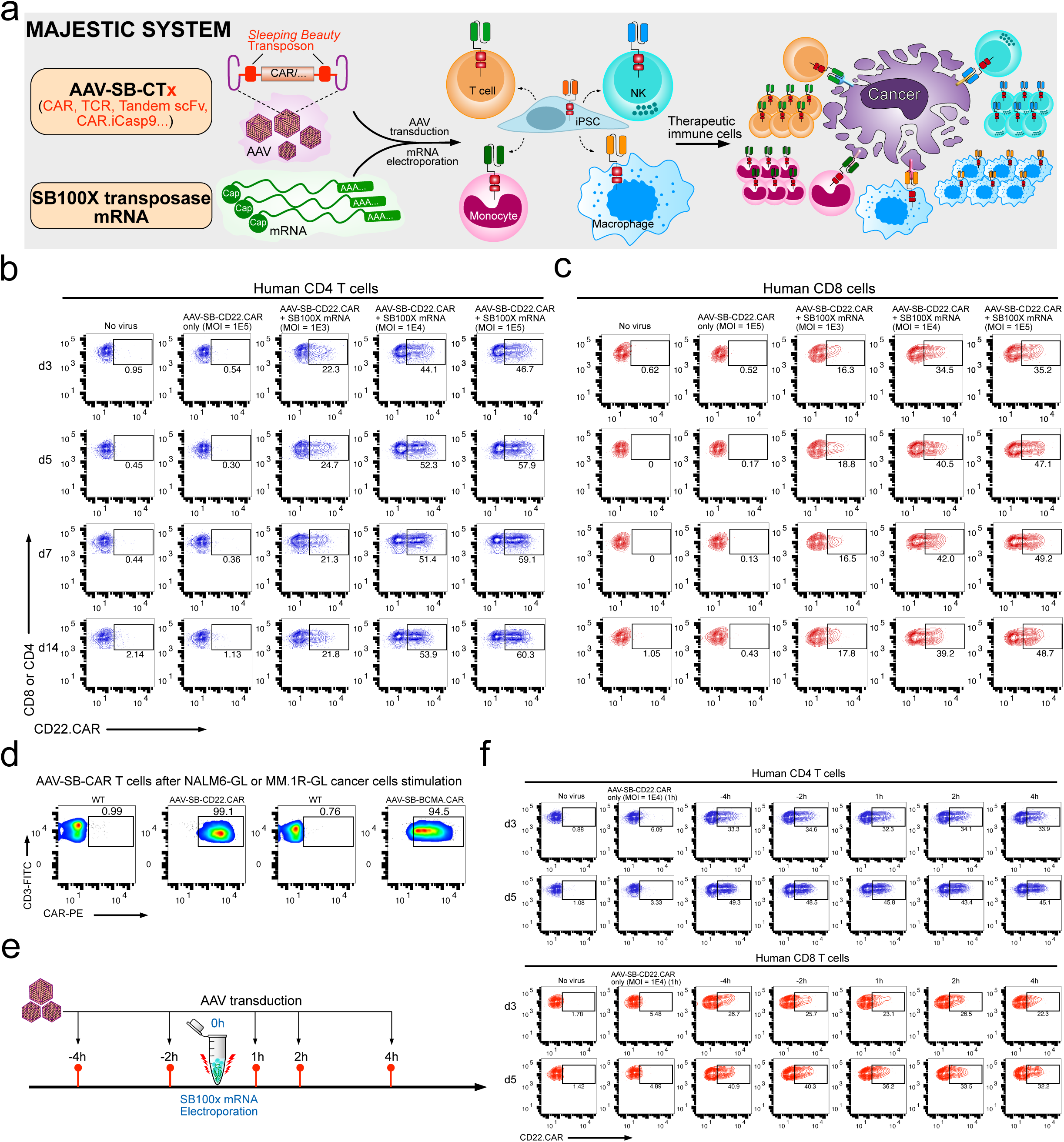
Development of MAJESTIC, a composite mRNA:AAV-SB system for highly efficient generation of therapeutic immune cells. **(a)** Schematics of the hybrid AAV-SB construct, SB100X mRNA electroporation, and CAR-T/NK/macrophage/iPSC generation. **(b)** Representative flow cytometry plots of human CD4 (gated for CD3^+^CD8^-^ cells) AAV-SB-CD22.CAR T cells. Human CD3 T cells were first electroporated with SB100x mRNA, then transduced with a titration series of AAV-SB-CD22.CAR virus. CAR-expression levels were evaluated at various time points from day 3 to day 14. **(c)** Representative flow cytometry plots of human CD8 (gated for CD3^+^CD8^+^ cells) AAV-SB-CD22.CAR T cells. Human CD3 T cells were first electroporated with SB100x mRNA, then transduced with a titration series of AAV-SB-CD22.CAR virus. Data were collected at several time points from day 3 to day 14. **(d)** Representative flow cytometry plots of AAV-SB-CD22.CAR and AAV-SB-BCMA.CAR T cells after cancer stimulation. **(e)** Schematic representation of various AAV-SB-CD22.CAR transduction time points relative to SB100x mRNA electroporation. **(f)** Representative flow cytometry plots of CD22.CAR T cells quantifying CAR-percentages of CD4 (top) and CD8 (bottom) T cells which were transduced with AAV-SB-CD22.CAR virus at various time points relative to when SB100x mRNA electroporation occurred (at 0h). In this figure, for optimization of conditions, each assay was done with one donor with three technical replicates. Donor 2 T cells were used in this figure. Cells were not purified for CD8/CD4 populations before electroporation.

We first examined the feasibility of this system for CAR-T cell generation. We electroporated human CD4 and CD8 T cells with SB100x mRNA, then transduced them with a titration series of AAV-SB-CD22.CAR virus **(Fig. S1a)**, using multiplicities of infection (MOIs) of 1E3, 1E4, and 1E5 **(Fig. 1b, c)**. We then monitored CD22.CAR expression via flow cytometry from day 3 to day 14. CD22.CAR-positive ratio correlated positively with virus titer, and CAR constructs were stably expressed in all time points tested. At day 14, for MOIs of 1E3, 1E4, and 1E5, respectively, 21.8%, 53.9%, and 60.3% of the total CD4 T cell population were CAR-positive **(Fig. 1b; Fig. S1c)**, and 17.8%, 39.2%, and 48.7% of CD8 T cells were CAR-positive **(Fig. 1c; Fig. S1c)**. We next tested a different CAR transgene, a BCMA-targeting CAR: at day 5 and at an MOI of 1E5, 35.6% and 32% of CD4 and CD8 T cells were BCMA.CAR-positive, respectively **(Fig. S1b, d)**. Furthermore, T cells in the AAV-SB-CAR condition can be quickly enriched for CAR-positive cells to nearly a pure CAR population following one-time antigen-specific cancer cell stimulation (day 12 post-stimulation), where CD22.CAR+ and BCMA.CAR+ T cells were 99.1% and 94.5% of cell populations, respectively **(Fig. 1d)**. These titration and condition-optimization experiments suggested that the MAJESTIC system can efficiently generate stable CAR-T cells from a single human donor.

Next, we surveyed the optimal time point(s) for AAV-SB-CTx viral transduction relative to SB100x mRNA electroporation. Setting SB100x mRNA electroporation as the 0h time point, we transduced cells with AAV-SB-CTx along a series of time points between -4h and 4h **(Fig. 1e)**. We observed that the AAV only group showed low CAR expression, although significant over the no-virus control background, potentially due to the transient expression of AAV **(Fig. 1f, g; Fig. S2a)**. In contrast, the MAJESTIC (AAV-SB-CTx) group has substantially higher CAR expression **(Fig. 1f, g; Fig. S2a)**. While the time points have similarly high CAR+ rates, the -4h time point (i.e. AAV-SB-CTx transduction 4h before mRNA-SB100x electroporation) appeared to be the most efficient numerically, which was true for both CD4 and CD8 T cells **(Fig. 1f, g; Fig. S2a, b)**. For practical convenience, we used a timepoint of 0h-1h for all experiments moving forward, because the efficiency differences between AAV transduction time points are moderate.

We then examined the optimal concentration of SB100X transposase. We titrated SB100X mRNA concentration by using a fixed amount of virus (MOI = 1E5) and varying amounts of mRNA **(Fig. 2a; Fig. S2c, d)**. Both CD4 and CD8 T cells electroporated with 1μg mRNA per 2e6 T cells yielded around 40% CAR-positive T cells. The CAR ratio is higher with 2μg as compared to 1μg mRNA in both CD4 and CD8 T cells; beyond 2μg of mRNA the CAR ratio appeared to be saturated **(Fig. 2a; Fig. S2c, d)**. We used a ratio of 2μg of mRNA per 2e6 cells hereafter.

**Figure 2.**
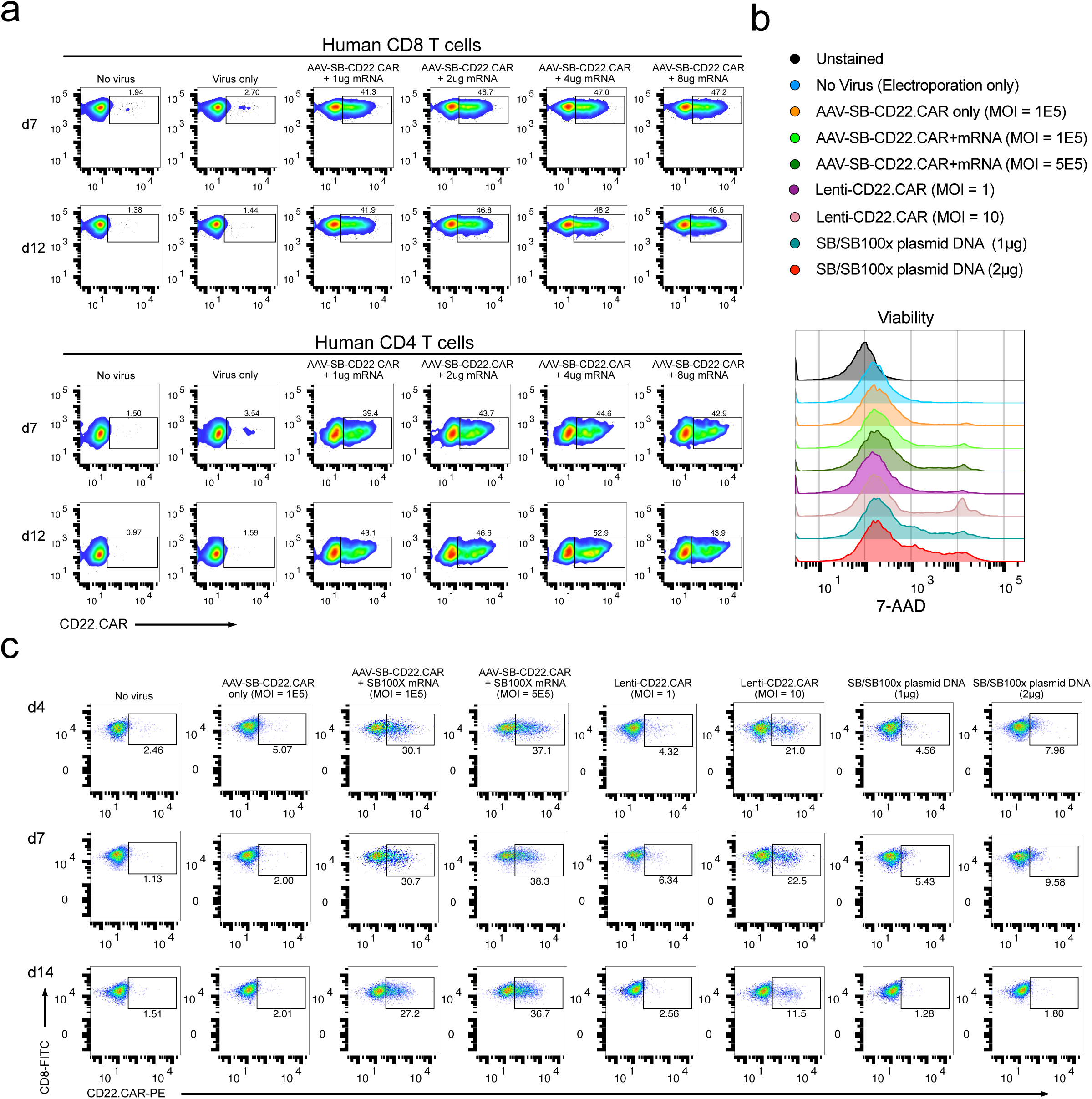
Comparison between MAJESTIC and conventional CAR-T generation approaches. **(a)** Representative flow cytometry plots of CD8 and CD4 CD22.CAR T cells were produced via AAV-SB and a titrated series of SB100X mRNA. **(b)** Representative flow cytometry plots measuring the viability of CD8 T cells. 7-AAD staining was performed to measure cell viability after mRNA electroporation, plasmid DNA electroporation, and lentivirus transduction. Media were changed one day after electroporation to remove the virus. **(c)** Representative flow cytometry plots of human CD8 T cells transduced with AAV-SB-CD22.CAR virus (MOI = 1E5 and 5E5), transduced with CD22.CAR lentivirus (MOI = 1 and 10), or electroporated with plasmid DNA (1μg = 0.5μg transposon plasmid + 0.5μg transposase plasmid) at three time points. In this figure, each assay was done with one donor with at least three technical replicates, donor 0286 T cells were used in this figure.

### MAJESTIC system efficiently produces stable, functional CAR-T cells with high viability and yield

We then compared CAR-T generation with the MAJESTIC system to lentiviral vector and SB transposon/transposase plasmid DNA electroporation approaches. It is difficult to uniformly compare these methods, as the underlying principles for each method differ. However, we still sought to systematically evaluate the performance of each method in parallel under standard laboratory settings (Methods). It is important to note that for AAV, the functional multiplicity of infection (MOI) (in transduction units) is usually 3-4 orders of magnitude lower than that of genomic MOI (in genome copies, GCs, gcs)^46^; therefore, in our experiments we used AAV genomic MOIs between 1E4 – 5E5. We used lentiviral MOIs between 1 – 10 according to current standard lentiviral CAR-T in literature ^47^. For DNA plasmid electroporation, we used between 1 - 4 μg total DNA / 1e6 cells, which is similar to the range of concentrations tested previously^38, 48^.

After CAR-T generation with all three platforms (MAJESTIC, lentivirus and SB transposon plasmid electroporation), we first determined cell viability via 7-AAD staining 48h after electroporation. Flow cytometry results showed that the MAJESTIC (AAV-SB-CD22.CAR + SB100x-mRNA) groups had viability above 85%. Lenti-CD22.CAR cells were over 90% viable at an MOI of 1 but only about 75% viable at a higher MOI of 10 **(Fig. 2b; Fig. S2e)**. Cells electroporated with plasmid DNA demonstrated lower viability: 75% and 70% for 1μg and 2μg of plasmid DNA, respectively **(Fig. 2b; Fig. S2e)**. Under these conditions, flow cytometry on day 4 revealed that Lenti-CD22.CAR at a high MOI (MOI = 10) yielded 21% CD22.CAR-positive CD8 T cells and only 4.32% at a low MOI (MOI = 1) **(Fig. 2c).** Even lower CAR-positive population percentages were observed for plasmid DNA electroporation groups (4.56% at 1μg and 7.96% at 2μg) compared to lentivirus **(Fig. 2c)**. Substantially higher CAR expression was observed in the MAJESTIC groups, with 30.1% (MOI = 1E5) and 37.1% (MOI = 5E5) CD22.CAR-positive T cells **(Fig. 2c)**. We proceeded to monitor the stability of CAR-positive T cell populations by examining CD22.CAR T populations at day 7 and day 14 by flow cytometry. CAR-positive populations were largely stable in the MAJESTIC groups, but significantly declined in the lentivirus and plasmid groups **(Fig. 2c; Fig. S2f)**.

We performed another experiment to specifically analyze the phenomenon that Lenti-CAR% declined across time in culture. After viral transduction and/or electroporation, we sorted Lenti-CAR and MAJESTIC-CAR T cells on day 2. Both normal Lenti-CD22.CAR and spin-infected Lenti-CD22.CAR groups demonstrated reduction of CAR+ percentages by day 5 (68.3% and 61.3%), falling even further by day 13 (52% and 43%). MAJESTIC-CAR T cells maintained a stable CAR+ ratio of around 85% (89.3% right after sorting) **(Fig. S2g)**. Although it is not entirely certain how lentiviral CAR ratios decline, it has been known that lentivirus transgenes can often get silenced ^22, 49^, potentially leading to reduced CAR percentages. CAR-T cell yield is an important joint outcome of viability, efficiency and proliferation (yield = CAR positive percentage * total live cell count), all of which can be affected by the cell states post CAR transgene delivery. We estimated the yield on day 5, 9, and 14. As a result, starting from approximately equal quantities of human CD8 T cells, the yields of CAR+ T cells from the MAJESTIC groups were much higher than those of lentivirus and of plasmid electroporation (e.g. 4.5x and 73.7x higher, comparing high dose conditions, respectively) **(Fig. S2h)**. These data suggested that, in the laboratory settings specified (see Methods), the MAJESTIC system is much more efficient in generating viable and stable CAR-T cells than lentiviral or DNA transposon electroporation approaches. Moreover, we also tested MAJESTIC using the Maxcyte electroporation platform: the procedure of MAJESTIC using both Neon and Maxcyte electroporators for introduction of the SB transposase (SB100x) mRNA component into cells yielded CAR percentages of over 36% **(Fig. S2i)**, suggesting that MAJESTIC is not limited to one electroporation platform.

### Initial characterization of CAR-T cells generated by MAJESTIC

We then sought to estimate the vector copy number (VCN) of CAR transgenes per cell in CAR-T cells generated by MAJESTIC, choosing to use a later time point (day 21) with the aim to measure stable transgenes. Using a standard approach similar to others in the field ^50^ (Methods), we estimated the VCN of MAJESTIC- and MC-SB + SB100X mRNA-generated CD22 CAR-T cells in four different human donors. The data showed that, under this experimental condition, the AAV-only group shows an average VCN of approximately 1. This detected background level could be due to prolonged episomal persistence of the AAV vector, and/or random genomic integration of the AAV vector DNA. MC-SB + SB100X mRNA group had a VCN of approximately 1-9 copies/cell. MAJESTIC group was observed with VCNs of approximately 1-4 copies / cell **(Fig. S3a-c)**. VCN measurement using both left arm and right arm probes showed consistent results **(Fig. S3a-c)**. We also quantified excision circles as a proxy for the excision efficiency of the SB100X transposase. AAV-SB-CAR constructs were significantly processed after one day of SB100X mRNA electroporation. We used qPCR primers amplifying the junction between the AAV arms and SB arms, for both the left and right sides separately to verify the assay. In amplifying the left arm, excision efficiency was around 55% on day 2 and 36% on day 3 **(Fig. S3d)**. With the right arm, excision efficiency was consistent, at around 53% on day 2, and 34% on day 3 **(Fig. S3d)**. Of note, the excision efficiency was slightly higher on day 2 compared to day 3, which may be due to the degradation of SB100X mRNA. This aligns the goal of using mRNA in MAJESTIC system in order to minimize unnecessary transposon excision after transgene delivery.

To address whether the CAR-T cells generated by MAJESTIC were functional, we first performed cancer killing assays by co-culturing cancer cells and CAR-T cells, generated either via MAJESTIC or lentivirus. We excluded the DNA plasmid electroporation group, as plasmid electroporated cells did not expand well and the yields were too low to be practically useful. Two forms of CAR were evaluated in co-culture: CD22.CAR vs. NALM6-GL cancer cells and BCMA.CAR vs. MM.1R-GL cancer cells. In counting T cells for kill assays, we normalized the number of T cells applied by CAR T ratio to ensure each group received the same number of CAR-T cells. While both MAJESTIC-generated CAR-T cells (AAV-SB-CAR) and lentiviral-transduced CAR-T cells (Lenti-CAR) exhibited tumor cell killing across the board, both AAV-SB-CD22.CAR and AAV-SB-BCMA.CAR T cells manifested significantly stronger killing over their lentivirus-mediated counterparts, for matched Effector : Target (E:T) ratios **(Fig. S3e, f)**. These data demonstrate that the CAR-T cells generated by MAJESTIC were indeed functional, with a potential advantage over the lentiviral CAR system in the settings tested.

We then further characterized the surface markers of MAJESTIC-generated CD8 CAR-T cells and explored whether virus and mRNA introduction would affect the immune phenotypes of T cells. We evaluated T cell exhaustion and memory markers before and after electroporation and viral transduction by staining for CD22.CAR and HER2.CAR T cells. The flow cytometry data showed that PD-1 was slightly decreased post MAJESTIC; while CTLA-4, TIM-3, and LAG-3 were increased; nevertheless, PD-1, CLTA-4, and TIM-3 all remained at baseline level as measured by mean fluorescence intensity (MFI) **(Fig. S3g)**. For the memory markers, only CCR7 demonstrated consistent increase in both HER2.CAR and CD22.CAR T cells. IL-7Ra remained at baseline for CD22.CAR and decreased for HER2.CAR **(Fig. S3g)**. CXCR3 demonstrated differing expression patterns, potentially due to differences in the CAR constructs (e.g., regarding costimulatory domains, CD22.CAR has 4-1BB, while HER2.CAR has 4-1BB and CD28). To further assess whether the CAR-T cells generated by MAJESTIC were effective against cancer, we performed a CAR-T efficacy testing experiment in vivo using an animal model of B cell leukemia with adoptive cell transfer treatment. Of note, this *in vivo* experiment was intended only to validate that MAJESTIC-generated CAR-T were indeed functional, but not to compare with cells generated by other platforms. The results showed strong anti-tumor efficacy, where MAJESTIC produced CAR-T cells, but not unmodified CD8 T cells, substantially suppressed cancer progression as measured by IVIS-bioluminescence imaging (p < 0.0001) **(Fig. S3h, i)**, and significantly extended the overall survival of treated mice (p = 0.0002) **(Fig. S3j)**. Of note, the animals in this cohort were solely used for efficacy study (IVIS imaging and survival) to demonstrate that MAJESTIC-produced CAR-T cells are functional, thus they were not euthanized concurrently. In the future, it will be feasible and informative to use MAJESTIC to generate CAR-T cells and evaluate their phenotypes such as persistence potential *in vivo*. Together, these data showed that the MAJESTIC system can efficiently produce stable and functional CAR-T cells with high viability and yield.

### Application of MAJESTIC in an array of different types of therapeutic transgenes in human T cells

We next applied this technology for engineering other types of therapeutic transgenes in human primary T cells, such as solid tumor CAR-T, tandem scFv (bispecific) CAR-T, TCR-T, and suicide-gene CAR-T cells **(Fig. 3a)**. For solid-tumor-specific CARs, we generated HER2-specific CAR-T cells, again comparing MAJESTIC, lentivirus, and plasmid DNA electroporation gene-transfer methods **(Fig. 3a, b; Fig. S4a)**. We observed that MAJESTIC produced the highest percentage of HER2.CAR-positive cells (28.1%) **(Fig. 3c; Fig. S4b)**. Lenti-HER2.CAR and plasmid DNA groups were less efficient, at 8.99% and 13.7%, respectively **(Fig. 3c; Fig. S4b)**. We also generated EGFRvIII-specific CAR-T cells in a similar manner. Although efficiency was generally lower, again, MAJESTIC’s CAR% T cell production efficiency was significantly higher than those of lentivirus **(Fig. S4c, d)**.

**Figure 3.**
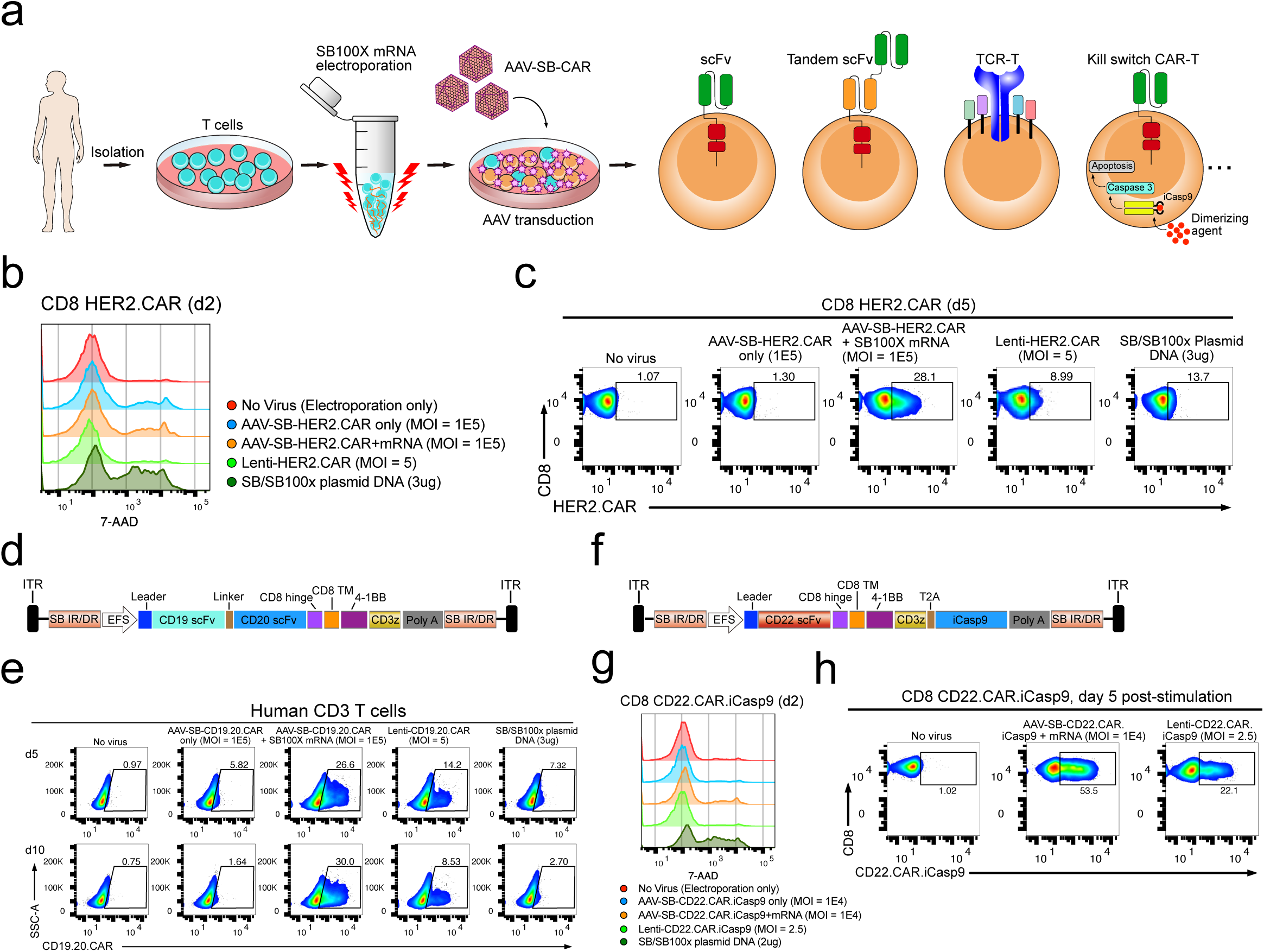
Application of MAJESTIC for delivery of different therapeutic transgenes into T human cells. **(a)** Schematic of various therapeutic cells (e.g. single scFv CAR, tandem scFv CAR, TCR-T, and kill-switch CAR) that can be generated via the AAV-SB system. **(b)** Flow cytometry and histogram overlays of cell viability post-electroporation as measured with 7-AAD staining. **(c)** Representative flow cytometry plots of HER2.CAR T cells. **(d)** Schematic representation of the AAV-SB-CD19.20.CAR construct. CD19 scFv and CD20 scFv CAR sequences are joined by a linker and are expressed together as a tandem scFv CAR. **(e)** Representative flow cytometry plots to evaluate CAR expression of CD19.20.CAR T cells. **(f)** Schematic representation of the AAV-SB-HER2.CAR.iCasp9 construct. **(g)** Flow cytometry and histogram overlays of cell viability post-electroporation as measured with 7-AAD staining. **(h)** Representative flow cytometry plots of CD22.CAR.iCasp9 T cells post antigen-specific cancer cells stimulation. In this figure, each assay was done with one donor with 3-5 technical replicates, donor 2 T cells were used for b, c; donor VP2 T cells were used for d, e; donor 2 T cells were used for g, h.

Bispecific CAR-Ts can recognize two antigens and may thereby reduce the chance of immune escape; such systems have demonstrated potent efficacy against relapsed B cell malignancies that down-regulated single target antigen expression ^20^. To test if MAJESTIC can be used for bi-specific CAR-T generation, we designed an anti-CD19/anti-CD20 tandem scFv construct, and used it to transduce primary human T cells, again comparing MAJESTIC to lentivirus and DNA transposon systems in parallel **(Fig. 3d)**. We performed flow cytometry on day 5 and day 10 after electroporation and viral transduction. As with single CARs, we observed that the efficiency of MAJESTIC was significantly higher than that of lentivirus, under the conditions tested **(Fig. 3e; Fig. S4e)**. Again, the CAR+% of the bispecific CAR-T cells was stable in the MAJESTIC group but was significantly reduced for both the lentivirus and DNA transposon systems **(Fig. 3e; Fig. S4e)**. In evaluating antigen expression of the cognate leukemia cancer cells, we noted high CD19 expression but weak CD20 expression **(Fig. S4f)**, consistent with previous reports^51^. The CD19.CD20 bispecific CAR-T cells exhibited strong killing ability, with nearly 100% and around 80% cytolysis after 17h at E:T ratios of 1:4 and 1:10, respectively, for both MAJESTIC and lentiviral CAR-T cells **(Fig. S4g)**. These data suggested that MAJESTIC can efficiently deliver a bispecific CAR construct into human T cells.

To test the utility of MAJESTIC for TCR-T cell production, we next cloned an NY-ESO-1 TCR construct along with a GFP marker into the AAV-SB backbone. AAV-SB-NY-ESO-1.GFP transduction plus SB100x mRNA electroporation was able to generate a fraction of NY-ESO-1 TCR-T cells, which was still significantly higher than that using SB/SB100x plasmid DNA electroporation (17% vs 11%) **(Fig. S5a, b)**. Conditional inactivation of CAR-T cells, e.g. through kill-switch elements such as induced Caspase 9 (iCasp9)^52^, may be clinically important for controlling potential toxicity. We thus generated conditional control CAR-T cells (CD22.CAR.iCasp9 T cells) with two transgenes, CD22 CAR and a suicide-gene **(Fig. 3f)**. Flow cytometry data showed that these cells were more efficiently generated by MAJESTIC than by lentiviral and plasmid systems under the experimental conditions as above **(Fig. 3g; S5c-e)**. CAR-positive T cells were further enriched in the AAV-SB-CD22.CAR.iCasp9 + mRNA group (53.5%) at day 5 after antigen-specific cancer cells stimulation **(Fig. 3h; Fig. S5f).** These data together demonstrated the versatility of the MAJESTIC system for delivering a variety of payloads to generate various therapeutic T cells.

### Head-to-head comparison of MAJESTIC system with mini-circle DNA system

The mini-circle (minicircle, MC) vector is a recently developed non-viral strategy that has shown significant improvement compared to conventional plasmid vector - specifically, the *SB* MC delivery^35^ of a CAR transgene has been proven to be more effective and less toxic compared with *SB* plasmid gene transfer. We performed a head-to-head comparison of the MAJESTIC system with both MC and plasmid DNA systems, using CD3 T cells as a source. 7-AAD staining data of SB/SB100X plasmid DNA and MC-SB/MC-SB100X groups showed similar cell viability (∼70%), with MC-SB + SB100X mRNA group showing slightly higher cell viability **(Fig. S6a)**. In comparison, AAV-SB-CD22.CAR + SB100X mRNA group (MAJESTIC) attained 86% viability, which was higher than SB/SB100X plasmid DNA, MC-SB/MC-SB100X, and MC-SB + SB100X mRNA groups **(Fig. S6a)**. CAR-T ratio on day 4 after electroporation confirmed higher CAR-T production efficiency compared of minicircle vs. plasmid DNA electroporation (19.7% vs. 8.01%). Efficiency could be further improved if MC-SB was electroporated with transposase supplied as SB100x mRNA (24.4%) **(Fig. S6b, c)**. In comparison, the MAJESTIC (AAV-SB-CD22.CAR + SB100X mRNA) group yielded the highest CAR-T ratio (51.8%) at day 4, substantially higher than those of MC/MC-transposase, MC/mRNA-transposase, and transposon plasmids **(Fig. S6b, c)**.

To further verify these results, we performed another independent set of MC vs MAJESTIC comparisons in an independent human donor, using CD4 and CD8 T cells separately in this case. Flow cytometry revealed CD22.CAR T ratios of 16.1%, 20.0%, and 24.2% for the MC-SB + SB100X mRNA group in donor 6760 CD4 T cells at day 3, 8, and 14, respectively **(Fig. 4a, c)**. The MAJESTIC group showed CD22.CAR T ratios of 43.4%, 60.9%, and 81.9% in donor 6760 CD4 T cells at day 3, 8, and 14, respectively **(Fig. 4a, c)**. Similar results were also observed in donor 6760 CD8 T cells. The MC-SB + SB100X mRNA group showed CD22.CAR T ratios of 29.1%, 29.4%, and 37.0% at day 3, 8, and 14, respectively **(Fig. 4b, d)**, and the AAV-SB-CD22.CAR + SB100X mRNA group showed CD22.CAR T ratios of 60.4%, 76.3%, and 82.4% at day 3, 8, and 14, respectively **(Fig. 4b, d)**. The yield of MAJESTIC and MC/mRNA-transposase is shown from aggregated replicates **(Fig. S6d)**.

**Figure 4.**
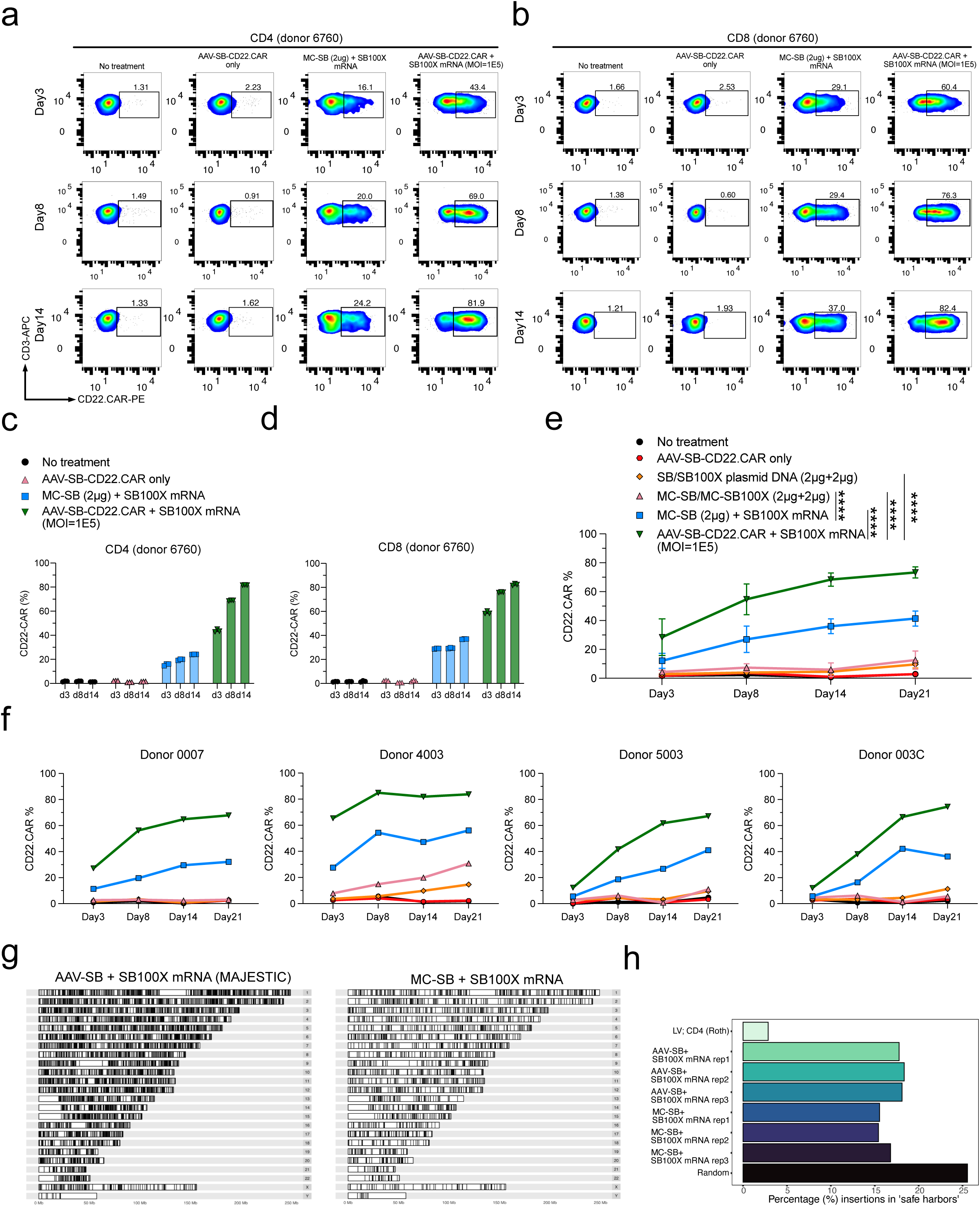
Comparison of minicircle transposon and MAJESTIC methods. **(a)** Flow cytometry of CD22.CAR ratio in donor 6760 CD4 T cells. **(b)** Flow cytometry of CD22.CAR ratio in donor 6760 CD8 T cells. **(c, d)** Quantification of flow cytometry data of CD22.CAR T cells from human primary CD4 (**c**) or CD8 (**d**) T cells, from an independent donor, with CAR-Ts produced by plasmid transposon plasmid, transposon MC, transposon MC with mRNA-transposase, and MAJESTIC (AAV-SB-CD22.CAR + SB100x mRNA). **(e-f)** Quantification of time-course flow cytometry data of CD22.CAR T cells produced by MAJESTIC and other systems, generated from PBMCs of four independent healthy human donors (n = 4) (**e**, showing mean ± s.e.m., n = 4 for all time points; **f**, showing individual donors in separate panels). **(g)** Karyogram of genomic insertions for one representative biological replicate from MAJESTIC and MC-SB + SB100X mRNA groups. **(h)** A barplot of frequencies of insertions in safe harbors. Three biological donors (n = 3) were mapped for MAJESTIC and MC-SB + SB100X mRNA groups. Random refers to a set of one million randomly generated genomic coordinates. Insertion coordinates for LV; CD4 (Roth) were obtained from literature (see main for citations). LV = lentivirus, MC = minicircle, SB100X = hyperactive Sleeping Beauty transposase. In this figure, data in a-d were sourced from one donor with three technical replicates; data in e, f were sourced from four independent donors (donor 0007, donor 4003, donor 5003, donor 003C). Two-way ANOVA with multiple comparisons tests was used to evaluate statistical significance.

To further test the variability of the MAJESTIC platform, we applied this technology along with other systems to four new healthy donors (Donor 0007, Donor 4003, Donor 5003, Donor 003C) in a head-to-head comparison manner. Consistently with the results above, across all four donors, the CAR-T ratio was the highest in MAJESTIC group (AAV-SB-CD22.CAR + SB100x mRNA) compared with other groups including MC/MC and MC/mRNA-Transposase **(Fig. 4e, f; Fig. S6e)**. Specifically, the CD22.CAR% was on average 29.20% ± 12.57% on day 3 (n = 4, mean ± s.e.m.), and as high as 73.3% ± 3.83% on day 21 for MAJESTIC (n = 4, mean ± s.e.m.) **(Fig. 4e; Fig. S6e)**. Importantly, although there is donor-to-donor variability as expected, in each respective donor, MAJESTIC is consistently the group with the highest efficiency in matched comparisons, across each donor and in all time points **(Fig. 4f)**. Electroporation involving either the plasmid or the MC form of DNA appears to have lower viability and CAR% vs. MAJESTIC even in different cell types and donors **(Fig. 4a-f; Fig. S6)**. Together, the data demonstrated the efficiency and reduced cellular toxicity of the MAJESTIC gene transfer platform compared with plasmid and MC gene transfer methods.

### Analyzing the genomic integration profile of the MAJESTIC system in CAR-T cells

To examine the genomic integration profile of the MAJESTIC system in T cells, we performed Splinkerette library preparation followed by next-generation sequencing (NGS) (Methods). We first purified CAR-positive T cells, then isolated genomic DNA from three sets (three independent donors) of MAJESTIC-generated and MC/mRNA-generated CAR-T samples collected d21 after electroporation. We fragmented the gDNA for these two donors and then conducted a two-step Splinkerette PCR to generate insertion site libraries. Analysis of next-generation sequencing data allowed us to map insertion locations in karyograms (**Fig. 4g)**. These data showed that MAJESTIC indeed mediates cargo integration into the genome of human T cells, across all major chromosomes.

We then examined the frequency of insertions into safe harbors, which are generally defined as regions of the genome where transgene insertions lead to predictable expression and do not interrupt existing gene activity^53^. We used a list of safe harbor coordinates from Querques et al. Using a random set of 1 million genomic sites, we estimated that approximately 25% of these random sites intersect with safe harbors. Using this value as a reference point, we compared MAJESTIC to other gene-transfer methods, using integration profile data from the literature for lentivirus transduction^54^. We sought to understand trends in the safety profile of MAJESTIC compared to other methods. While MC/mRNA-mediated insertions were similar to MAJESTIC to be within safe harbors (around 15% vs. 17% on average), MAJESTIC-mediated insertions were much more likely to be within safe harbors compared to that of lentivirus (around 2∼3%) (**Fig. 4h)**. Furthermore, we determined the proportion of insertions into functional gene regions, including exons, introns, and cancer genes and calculated the frequencies as fold-change relative to the randomly generated sites (**Fig. S7**). Using this functional-gene region profile as a proxy for insertion safety, MAJESTIC-insertions occupy a reasonably favorable performance, comparable to MC/mRNA and better than lentiviral vector (**Fig. 4g, h; Fig. S7a**). Together, these data reveal the integration site profile of the MAJESTIC system and demonstrate a trend towards safer insertions compared to lentiviral transduction.

Unexpectedly, we noticed certain differences in the frequencies of integration into exons, coding exons, and 5’ UTRs in MC-SB + SB100X mRNA group, as compared with a previous study^38^. This may be due to technical reasons (e.g., different sample preparation, different time point). For example, it is possible that the selection of the CAR+ cell population in this analysis might introduce a difference (because CAR+ selection enriches T cells with the CAR transgene inserted in genomic loci that avoid transgene silencing), as compared to the unselected bulk samples in the previous study ^38^. Regarding the integration profile differences observed between MAJESTIC and MC systems, because both have the same SB transposon, the differences could be due to technical (e.g. sample prep) and/or biological reasons associated with the differences in transposon delivery approaches (e.g. the ssDNA SB template provided by AAV in episome, and the dsDNA SB template provided by MC in transient extrachromosomal DNA).

### Application of MAJESTIC across multiple human immune cell types

While CAR-Ts have strong clinical potential, they also have inherent limitations^4^. It is well-known that the suppressive tumor microenvironment of solid tumors creates significant hurdles for T cells^55^. However, unlike T cells, myeloid cells such as macrophages and monocytes naturally infiltrate tumors ^56^. In addition, natural killer (NK) cells have been explored as an alternative to T cells for immunotherapy, because they utilize a different set of signaling pathways, have rapid activation, can exhibit TCR-independent cytotoxicity, and are relatively easier to develop into an off-the-shelf product^57^. Therefore, it is of interest to expand cell therapy to other immune cell types, such as NK cells and myeloid cells ^58^, to overcome the inherent limitations of T cell based therapy.

To test the utility of MAJESTIC in other immune cell types, we used MAJESTIC for delivery of CAR transgenes to generate CAR-NKs, CAR-Monocytes (CAR-Monos) and CAR-Macrophages (CAR-Mas). We again performed lentivirus transduction, and/or plasmid electroporation in parallel. NK92 is an immortalized NK cell line that has been used to produce CAR-NKs, which have achieved use in clinical trials^59^. We transduced NK92 cells to engineer HER2-specific CAR-NK cells. Flow cytometry revealed that the MAJESTIC system efficiently generated HER2 CAR-NK cells (nearly 50% at a high dose on day 14) at significantly higher rates than that by lentiviral or DNA transposon electroporation in the conditions tested **(Fig. 5a-b)**. For example, at day 4, Lenti-HER2.CAR at a high MOI (MOI = 2.5) yielded 13.9% HER2.CAR-positive NK92 cells, but only 2.71% with low MOI (MOI = 1) **(Fig. 5a-b).** Plasmid DNA transposon electroporation groups achieved 16.1% and 8.06% HER2.CAR-positive NK92 cells **(Fig. 5a)**. AAV-SB-HER2.CAR+mRNA groups yielded 41.6% and 13.2% HER2.CAR-positive NK92 cells in high (1E5) and low (1E4) MOI conditions, respectively **(Fig. 5a)**. Notably, the proportion of the HER2.CAR-positive population did not substantially decline in long-term cultures of MAJESTIC-generated CAR-NK cells: the percentage of HER2.CAR+% NK cells was 44.9-49.0% at day 14 **(Fig. 5a, b)** and was sustained at 43.8% on day 33 **(Fig. 5c, d)**. Of note, the dose-dependence effect is strong for CAR-NK generation via MAJESTIC, as high (1E5) MOI resulted in significantly higher CAR+% NK cells than low (1E4) MOI **(Fig. 5a-d)**. Because NK cells can kill cancer cells independent of cancer-specific antigens, we tested the function of the resultant HER2.CAR-NK cells in a kill assay with co-culture of MCF-7 breast cancer cells. Results showed that the HER2.CAR-NK cells are efficient killer cells, demonstrating significantly stronger killing compared to untreated NK92 cells, with around 70% target cell death 24h post co-culture **(Fig. 5e)**. These data demonstrate that the MAJESTIC system can be used to efficiently generate stable and functional CAR-NK cells.

**Figure 5.**
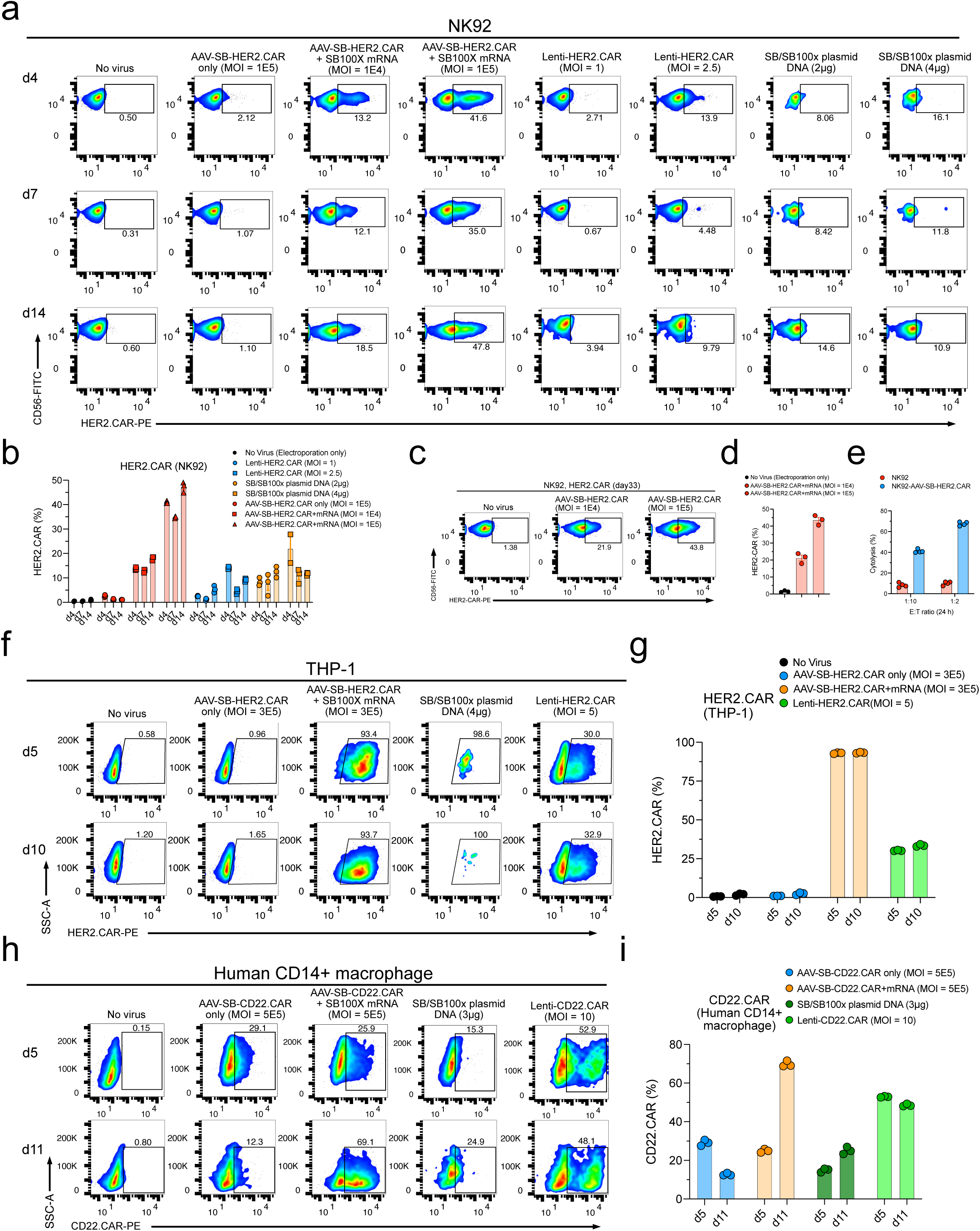
CAR-NK, CAR-Monocyte, and CAR-Macrophage generation via MAJESTIC. **(a)** Representative flow cytometry plots of NK92 cells transduced with AAV-SB-HER2.CAR virus (MOI = 1E4 and 1E5), transduced with HER2.CAR lentivirus (MOI = 1 and 2.5), or electroporated with plasmid DNA (2μg = 1μg transposon plasmid + 1μg transposase plasmid) at three different time points. **(b)** Quantification of (a). **(c)** Flow cytometry measurement of HER2.CAR expression in NK92 cells on day 33 after mRNA electroporation and AAV-SB-HER2.CAR viral transduction. **(d)** Quantification of (c). **(e)** Cytolysis analysis of MCF7-PL (MCF7 with puromycin resistance and luciferase expression) cancer cells that were co-cultured with NK92-AAV-SB-HER2.CAR cells. CAR-NKs were seeded at two effector : target (E:T) ratios, and a luciferase assay was performed at two time points (24h and 48h). **(f)** Representative flow cytometry plots of THP-1 cells (human monocytic cell line) transduced with AAV-SB-HER2.CAR virus (MOI = 3E5), transduced with HER2.CAR lentivirus (MOI = 5), or electroporated with plasmid DNA (4μg total). **(g)** Quantification of (f). **(h)** Representative flow cytometry plots of human CD14+ macrophages transduced with AAV-SB-CD22.CAR virus (MOI = 5E5), transduced with CD22.CAR lentivirus (MOI = 10), or electroporated with plasmid DNA (3μg). **(i)** Quantification of (h). In this figure, each assay has at least three technical replicates, and two-way ANOVA with multiple comparisons test and multiple unpaired *t* tests (with adjusted p value) were performed to evaluate statistical significance.

Next, we tested this system for CAR delivery to THP-1 cells, a human monocytic and myeloid cell line ^60^. We achieved over 93% HER2.CAR-positive THP-1 with AAV-SB-HER2 (MOI 3E5) on day 5 and 10, compared to ∼30% with lentiviral transduction at MOI 5 **(Fig. 5f, g)**. For the DNA transposon plasmid electroporation group, cell viability was extremely low, with the vast majority of cells (>99%) dead due to the high cellular toxicity of plasmid electroporation **(Fig. 5f, g)**, making it impossible to generate sufficient cells for subsequent analysis. We then set out to engineered CAR-MAs using primary human CD14^+^ macrophages, again comparing MAJESTIC with lentivirus and DNA transposon systems. Flow cytometry revealed a 25.5% CD22-specific CAR-MA population for the AAV-SB-CD22.CAR+mRNA group, which increased to 69.1% by day 11 **(Fig. 5h, i)**. Lentiviral transduction worked reasonably well in primary macrophages, with CAR-positive percentages of 52.9% on day 5 and 48.1% on day 11 **(Fig. 5h, i)**. Interestingly, AAV-SB-CAR alone without the mRNA-transposase yielded nearly 30% CAR+ MAs by day 5, which fell by more than half by day 11 **(Fig. 5h, i)**. At day 11, the CAR+% of CAR-MAs in the MAJESTIC group was highest among all groups tested (vs. AAV-SB-CAR alone, lentivirus and DNA transposon). These data suggest that the MAJESTIC system can be used to efficiently generate stable CAR-MA cells. Finally, because human iPSCs can be used as a source for derivation of various types of cellular lineages ^15 17, 18^, we set out to test if MAJESTIC can be applied to iPSCs. Results showed MAJESTIC can transduce iPSCs carrying a HER2.CAR transgene at high efficiency (>75% HER2.CAR+) **(Fig. S7b, c)**. Along with MAJESTIC, lentiviral vector can also transduce iPSCs at high efficiency, but not transposon DNA electroporation **(Fig. S7b, c)**. Although differentiation of iPSCs into other cell types takes additional time ^15 17, 18^, delivery of cell therapy transgenes into iPSCs by MAJESTIC provides another avenue to generate various therapeutic immune cells.

Altogether, these data suggested that the MAJESTIC system is capable of efficiently engineering stable functional therapeutic immune cells and is applicable to various types of transgenes and across multiple lineages of immune cells.

## Discussion

Adoptive cell therapy, most notably CAR-T therapy, has demonstrated clinical success in patients with several indications of hematological malignancies. A vital and potentially limiting step of this therapy is the manufacturing of engineered immune cells: to produce sufficient therapeutic cells for therapy would ideally require 1) a sizeable pool of patient immune cells to begin with and 2) a highly efficient genetic engineering technology. Although low transfer efficiency can be alleviated in part by increased culturing time, high initial efficiency would expedite the manufacturing process. Additionally, increasing culturing time may necessitate extended exposure of the cells to their target antigens during production, which may shift cells in favor of a differentiated phenotype, reducing long-term memory function ^21^ and thus the overall quality of the engineered cells.

γ-retroviral vectors are indeed capable of efficient genome integration, but their tendency to insert into promoters of actively transcribed genes raises concerns about potential genotoxicity. Lentiviruses are commonly used in clinical trials today, and recent advances have improved their safety. However, there are still certain safety risks associated with the pathogenic origin of such viruses, and a recent study has suggested a higher preference of lentiviral insertion into active genes compared to Sleeping Beauty-based gene transfer methods^61, 62^.

The field has developed various alternative gene transfer methods that can be worked with at BSL-1^21^. AAV is a commonly used gene therapy vector, however, due to the dilution effect, AAV-transduced T cells have a gradual reduction in transgene expression, making an AAV-only system not ideal for delivery of CAR transgenes into T cells. Electroporation of 1) DNA transposon/transposase or 2) CAR-encoding mRNA construct are two non-viral gene transfer strategies, but key limitations include low viability/high toxicity for the former, and transient transgene expression for the latter.

The notable success of immune cell-based cell therapies has invited the introduction of other technologies to enhance immune cell engineering. CRISPR is one such technology that is being rapidly and broadly applied to immune cell editing ^29, 30, 63^. Such strategies rely on Cas9, Cas12a/Cpf1 or other DNA-targeting endonucleases to generate DSBs, which are then repaired with a donor template via homology-directed repair (HDR). The efficiency of these knock-in/knock-out systems is thus dependent on two steps: 1) the efficiency of enzymatic gene knockout and 2) the rate of incorporation of the homology template. As to 1), knockout efficiency depends on the availability of an optimal guide because poorly designed guides may not cut efficiently and may cause undesired off-target effects ^64–69^. In addition, CRISPR knockout generates exposed DSBs that may trigger mutagenic non-homologous end-joining (NHEJ) pathways rather than HDR, which is especially hazardous for off-target editing, when no repair template is available for HDR to occur. As to 2), HDR is limited to the late S and G2 phases of the cell cycle^70^, restricting the interval in which the second step can occur. In CRISPR-based gene editing approaches, DSB occurs and brings two types of risks – the triggering of the p53 pathway, and the possibility of chromosome alterations, which increases with the number of DSBs^71^, for example, the observed chromosomal aberrations following Cas9-mediated genome editing in recent studies^31–33^. MAJESTIC itself does not involve CRISPR-based gene editing.

Unlike γ-retroviruses, SB reduces the likelihood of genotoxicity as studies have shown that this class of transposons has close to a random genomic integration profile ^72^. However, introduction of the SB system into cells by DNA transfection or electroporation can lead to higher cellular toxicity^38, 73^. During remobilization, SB can leave behind a tri-nucleotide footprint; thus continuous remobilization of the transposon is a potential limitation of an all-in-one AAV-SB system^44^, where both the transposon and transposase are delivered by AAV. Our platform addresses this issue by separating the transposase into a transient delivery component (mRNA). A prior study used a hyperactive transposase SB100X to improve CAR-T transduction of DNA transposon ^74^. The SB system has also been engineered in the form of combinations or hybrid vectors, e.g., dCas9-SB100X to retarget SB transposition^75^, and an adenovirus-SB hybrid system to achieve higher transduction efficiency^76^. The MAJESTIC system differs from all such efforts by combining the advantages of all three delivery vehicles (AAV, transposon, mRNA) in an organic way: transducing cells with the hybrid AAV-transposon vector with electroporation of transposase mRNA. AAV-SB transduction retains the benefits of high cell viability and stable transgene expression. The process of gene transfer of the MAJESTIC system is similar to conventional SB nucleofection, with mRNA electroporation instead of plasmid or MC electroporation and an extra AAV transduction step in which virus is added directly into the media. From our data, it appears that approaches involving DNA electroporation including plasmid transposon, MC/MC and/or MC/mRNA, naturally have an associated impact on cell viability and yield. MAJESTIC avoids introducing DNA into cells and instead uses AAV and mRNA, both of which have reasonably low cellular toxicity.

The limitations of the MAJESTIC system must be acknowledged. By virtue of relying on AAV for gene-transfer, the MAJESTIC system will be limited by AAV’s packaging size of ∼4.75kb (∼4.3kb without SB arms) (**Table S1**). This is usually sufficient to include the CAR construct and additional elements (e.g., iCasp9), but will face challenges with significantly larger transgenes, which could be accommodated with DNA transposon plasmid/MC systems as transposons can in principle carry large transgene cargos although the efficiency may drop as the size increases ^77^. Additionally, given that MAJESTIC is a composite system, the generation of therapeutic immune cells is a two-step process including electroporation/nucleofection + viral transduction, although they can be streamlined to be performed at the same period of time (as demonstrated in our 0h transduction/electroporation experiments); while lentivirus or plasmid electroporation are both one-step methods. Also, compared to lentivirus or plasmid production, the good manufacturing practice (GMP) production cost of MAJESTIC will be higher because of the requirement for both AAV and mRNA. Without considering yield, MAJESTIC, KIKO, AAV and lentiviral/retroviral approaches all have higher GMP cost as compared to non-viral approaches such as transposon/MC, which is more economic to manufacture per today’s GMP landscape (**Table 1**). MAJESTIC itself can not achieve precisely targeted gene editing as CRISPR, rather, its advantage is being a high-efficiency gene-editing-free delivery approach. We present MAJESTIC as an alternative cargo delivery and therapeutic cell generation strategy with the strength of producing CAR+ cells with high viability at high yield - and thus propose that the MAJESTIC system may be worth considering where viability and/or yield is particularly important.

**Table 1.**
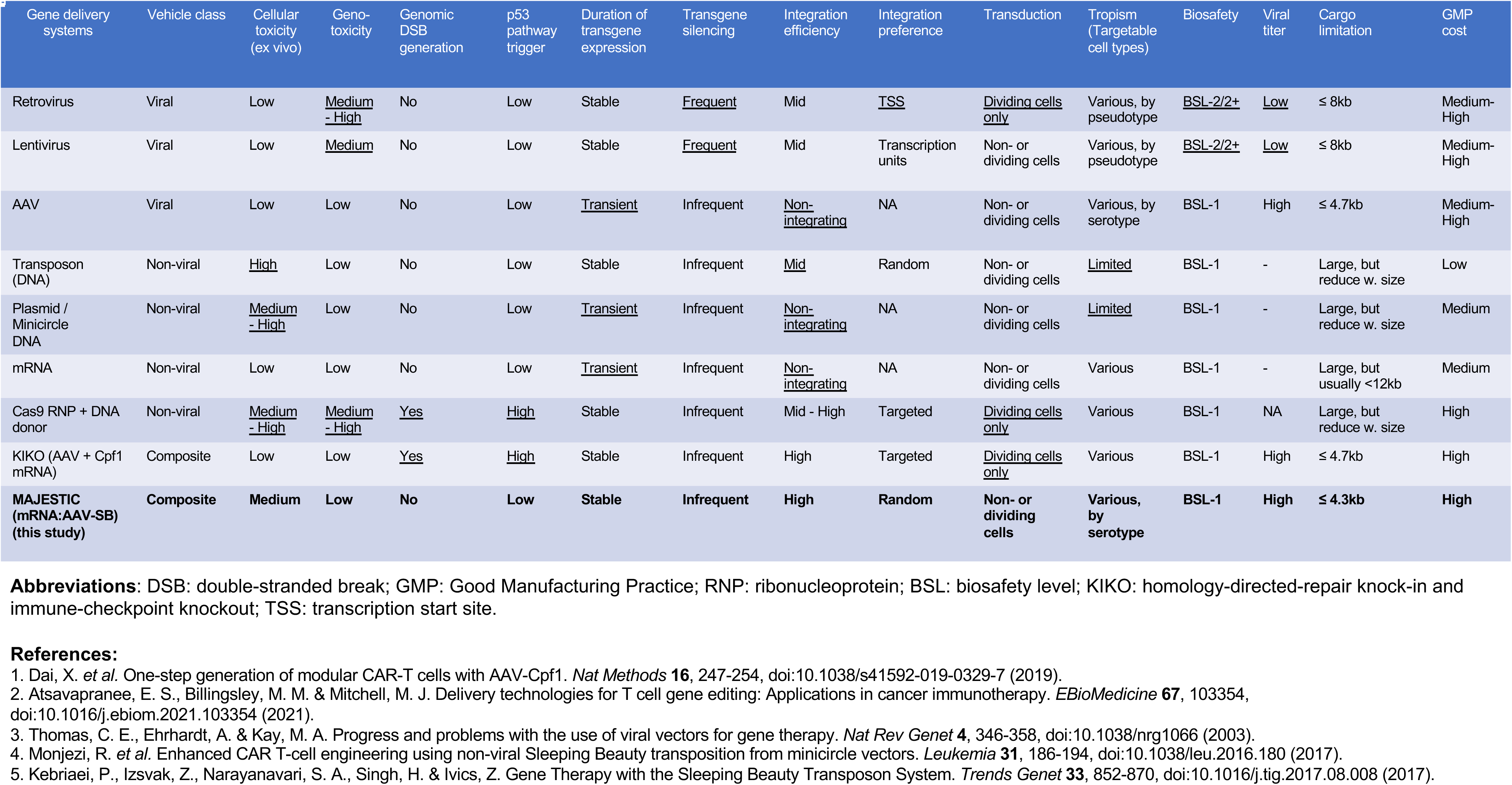
Comparison of gene delivery and cell therapy engineering technologies.

| Gene delivery systems | Vehicle class | Cellular toxicity (ex vivo) | Geno-toxicity | Genomic DSB generation | p53 pathway trigger | Duration of transgene expression | Transgene silencing | Integration efficiency | Integration preference | Transduction | Tropism (Targetable cell types) | Biosafety | Viral titer | Cargo limitation | GMP cost |
| --- | --- | --- | --- | --- | --- | --- | --- | --- | --- | --- | --- | --- | --- | --- | --- |
| Retrovirus | Viral | Low | <u>Medium - High</u> | No | Low | Stable | <u>Frequent</u> | Mid | <u>TSS</u> | <u>Dividing cells only</u> | Various, by pseudotype | <u>BSL-2/2+</u> | <u>Low</u> | ≤ 8kb | Medium-High |
| Lentivirus | Viral | Low | <u>Medium</u> | No | Low | Stable | <u>Frequent</u> | Mid | Transcription units | Non- or dividing cells | Various, by pseudotype | <u>BSL-2/2+</u> | <u>Low</u> | ≤ 8kb | Medium-High |
| AAV | Viral | Low | Low | No | Low | <u>Transient</u> | Infrequent | <u>Non-integrating</u> | NA | Non- or dividing cells | Various, by serotype | BSL-1 | High | ≤ 4.7kb | Medium-High |
| Transposon (DNA) | Non-viral | <u>High</u> | Low | No | Low | Stable | Infrequent | <u>Mid</u> | Random | Non- or dividing cells | <u>Limited</u> | BSL-1 | - | Large, but reduce w. size | Low |
| Plasmid / Minicircle DNA | Non-viral | <u>Medium - High</u> | Low | No | Low | <u>Transient</u> | Infrequent | <u>Non-integrating</u> | NA | Non- or dividing cells | <u>Limited</u> | BSL-1 | - | Large, but reduce w. size | Medium |
| mRNA | Non-viral | Low | Low | No | Low | <u>Transient</u> | Infrequent | <u>Non-integrating</u> | NA | Non- or dividing cells | Various | BSL-1 | - | Large, but usually <12kb | Medium |
| Cas9 RNP + DNA donor | Non-viral | <u>Medium - High</u> | <u>Medium - High</u> | <u>Yes</u> | <u>High</u> | Stable | Infrequent | Mid - High | Targeted | <u>Dividing cells only</u> | Various | BSL-1 | NA | Large, but reduce w. size | High |
| KIKO (AAV + Cpf1 mRNA) | Composite | Low | Low | <u>Yes</u> | <u>High</u> | Stable | Infrequent | High | Targeted | <u>Dividing cells only</u> | Various | BSL-1 | High | ≤ 4.7kb | High |
| **MAJESTIC (mRNA:AAV-SB) (this study)** | **Composite** | **Medium** | **Low** | **No** | **Low** | **Stable** | **Infrequent** | **High** | **Random** | **Non- or dividing cells** | **Various, by serotype** | **BSL-1** | **High** | **≤ 4.3kb** | **High** |

**Abbreviations:** DSB: double-stranded break; GMP: Good Manufacturing Practice; RNP: ribonucleoprotein; BSL: biosafety level; KIKO: homology-directed-repair knock-in and immune-checkpoint knockout; TSS: transcription start site.

We summarize the differences, including the advantages and limitations of the MAJESTIC system and compare it to existing approaches for therapeutic cell engineering in **Table 1**. It should be noted that MAJESTIC does not replace nor diminish other methods, instead, it provides a recently developed alternative gene delivery technology that offers superiority and advantages in certain feature areas, such as high viability, efficiency and yield; while naturally associated with limitations such as cargo size, additional procedure, and cost. The versatility of the MAJESTIC system makes it not limited to application in cell therapy for cancer - any therapy or research effort utilizing engineered immune cells could, in principle, benefit by using this system. Future research could examine the efficacy of MAJESTIC in generating CAR-Ts from cancer patient-derived T cells in translational studies to validate the utility in a more clinically relevant scenario. Further improvements in the safety of the platform can be achieved, using other components including different AAV serotypes, other viral vectors, or different transposon systems such as a high-soluble Sleeping Beauty transposase (hsSB) ^62^. Given the modular nature of the MAJESTIC platform, other cargos and cell types can be tested in a broad range of applications by users from various fields.

## Acknowledgments

We thank all members of Chen laboratory and various Yale entities for various discussions. We thank various Yale core facilities for technical supports. In particular, we thank K. Tang and P. Renauer for technical assistance on Illumina sequencing and data analysis.

S.C. is supported by Yale SBI/Genetics Startup Fund, NIH/NCI/NIDA (DP2CA238295, R01CA231112, R33CA225498, RF1DA048811), DoD (W81XWH-20-1-0072, W81XWH-21-10514), Alliance for Cancer Gene Therapy, Sontag Foundation (DSA), Pershing Square Sohn Cancer Research Alliance, Yale Cancer Center Pilot Award, Dexter Lu Gift, Ludwig Family Foundation, and Chenevert Family Foundation. S.Z.L is supported by Yale College Fellowships.

## Author Contributions

SC conceived the study. L. Ye designed the experiments with assistance from SL. L. Ye performed most experiments, with assistance from SL, LP, KS, QL, YZ and PC. L. Yang, KS and YZ performed 2^nd^ revision experiments with technical supervision by LP. SL and KS analyzed NGS data. L. Ye, SL, L. Yang and SC prepared the manuscript with inputs from all authors. SC secured funding and supervised the work.

## Methods

### Institutional Approval

This study has received institutional regulatory approval. All recombinant DNA and biosafety work were performed under the guidelines of Yale Environment, Health and Safety (EHS) Committee with approved protocols (Chen-rDNA-15-45; Chen-rDNA-18-45). All human sample work was performed under the guidelines of Yale University Institutional Review Board (IRB) with an approved protocol (HIC#2000020784). All animal work was performed under the guidelines of Yale University Institutional Animal Care and Use Committee (IACUC) with approved protocols (Chen-2018-20068; Chen-2021-20068).

### Mouse model

Mice were housed in standard conditions in Yale vivarium, maintained on a 12h light/dark cycle (07:00 to 19:00 light on). Mice, both female and male, aged 8-12 weeks were used for experiments. NOD-scid IL2Rgammanull (NSG) mice were purchased from JAX and bred in-house for T cell-based anti-tumor therapeutic efficacy testing experiments. Mouse health was monitored daily after tumor induction.

### Construction of AAV-SB-CAR vector

To create hybrid AAV-SB-CAR vectors, we began with our previously established AAV-SB-CRISPR vector as a backbone, which has an sgRNA/SB100x expression cassette nested between SB arms and AAV ITRs ^44^. We replaced this expression cassette between the U6 promoter and the short polyA sequence with a CAR or other expression cassettes. As this study utilized multiple types of CARs (e.g. CD22, BCMA), we obtained each CAR sequence via either 1) PCR amplification of CAR sequences from existing CAR constructs ^78^, or 2) IDT gene synthesis. The SB100x transposase was subcloned from ^44^ and cloned into the *Nco* I and *Hind* III restriction endonuclease sites of the empty vector pcDNA3.1, which was used for *in vitro* transcription of mRNA.

### Preparation of minicircle (MC) DNA

Genes-of-interest such as SB-CAR and SB100X constructs were firstly cloned into a parental plasmid (System Biosciences). Then MC DNA was produced and purified by using an MC-Esay^TM^ Kit (System Biosciences), without the optional dNTP removal step.

### Cell culture

HEK293T, NALM6, MM.1R, MCF7, NK-92, THP-1, human CD14+ monocytes, human PBMCs, and human iPSCs were purchased from commercial sources (ThermoFisher, American Type Culture Collection (ATCC), and STEMCELL). HEK293T and MCF7 cells were cultured in DMEM (Gibco) media supplemented with 10 % FBS (CORNING) and 200 U / mL penicillin– streptomycin (Gibco), hereafter referred to as D10. NALM6 and MM.1R cells were cultured in RPMI-1640 (Gibco) media supplemented with 10% FBS and 200 U / mL penicillin–streptomycin.

NK-92 cells were cultured in Alpha Minimum Essential medium (MEM) (Gibco) supplemented with 12.5% horse serum, 12.5% FBS, 0.2 mM inositol, 0.1mM 2-mercaptoethanol, 0.02mM folic acid, and 200U/mL human IL-2 (BioLegend).

THP-1 and CD14+ monocytes were cultured in RPMI-1640 media supplemented with 10% FBS, 1% Glutamax, 1% HEPES, and 1% penicillin–streptomycin. 20ng/mL of human GM-CSF (BioLegend) was used to differentiate monocytes into macrophages for 7 days, then macrophages were collected for electroporation and/or viral transduction, CAR-macrophage cells were maintained in RPMI complete media supplied with 20ng/mL of human GM-CSF.

Human PBMCs, CD4, and CD8 T cells were purchased from the StemCell and cultured in X-VIVO^TM^ 15 media (Lonza) supplied with 5 % human AB serum (MP Biomedicals) and 400U/mL human IL-2 (BioLegend). T cells were activated with Dynabeads Human T-Activator CD3/CD28 (ThermoFisher) with a T cell:Beads ratio at 1:1. In this study, multiple T cell donors were involved in various experiments; the donors for each experiment are clarified in figure legends.

Human iPSCs were cultured in StemFlex^TM^ Medium (Gibco).

### Lentivirus production and titration

Low-passage (less than 15 passages) HEK293T cells were used for lentiviral packaging. One day before transfection, 2e7 HEK293T cells were seeded per 150mm-dish. D10 media was replaced with 13 mL pre-warmed Opti-MEM medium (Invitrogen) before transfection. For each plate, 20μg transgene plasmid, 15μg psPAX2 (Addgene), 10μg pMD2.G (Addgene) and 90μL lipofectamine 2000 (Thermo Fisher) were mixed in 450μL Opti-MEM. The mixture was vortexed briefly and incubated for 10-15min at room temperature, then added dropwise to cells. To minimize the toxicity of lipofectamine, Opti-MEM media was replaced with pre-warmed 20mL D10 media 5-6 h after transfection. Viral supernatant was collected 48h post-transfection and then concentrated using the Amicon Ultra-15 Centrifugal filter unit (Millipore), or purified with Lenti-X Concentrator (Takara). All virus was titrated with Lenti-X GoStix Plus (Takara) before being aliquoted and stored in −80 °C.

### AAV production, purification, and titration

HEK293T cells were prepared in 150mm-dishes as above. D10 media was replaced by 13mL pre-warmed DMEM (FBS-free). For each 150mm-dish, HEK293T cells were transiently transfected with 5.2μg transfer, 8.9μg AAV6 serotype and 10.4μg pDF6 plasmids, which was pre-mixed with 130 μL of PEI (1 mg/mL) in 450 μL Opti-MEM medium. After 6h of transfection, DMEM was replaced with 20 mL pre-warmed D10 media. Transfected cells were dislodged and collected in 50mL Falcon tubes 72h post-transfection for AAV purification. AAV purification was performed as previously reported ^44^. Viral titer was measured via RT-qPCR with a Taqman probe targeting the EFS sequence in the AAV vector.

### Copy number determination

T cells were collected at d21 after electroporation and washed with PBS twice to remove media. Cells were incubated with CD22-Fc protein (R&D Systems) in PBS for 30 min on ice and washed with PBS twice to remove the unbonded protein. The cells were then stained with anti-human IgG Fc-APC (Biolegend, Cat#366906) on ice for 30 min and washed with PBS twice. The CAR-positive T cells were purified by Anti-APC MicroBeads (Miltenyi). After that, genomic DNA was extracted using the QIAGEN Blood Mini Kit. To minimize episomal AAV contamination, the extracted genomic DNA samples were separated on a 1% agarose gel, and DNA bands over 10kbp were gel isolated and purified with QIAquick Gel Extraction Kit (QIAGEN). We conducted qPCR to determine copy number, using primers that specifically target SB transposon:

IRDR-left (Forward: 5’-CTCGTTTTTCAACTACTCCACAAATTTCT-3’

Reverse: 5’-GTGTCATGCACAAAGTAGATGTCCTA-3’)

IRDR-right (Forward: 5’-GCTGAAATGAATCATTCTCTCTACTATTATTCTGA-3’

Reverse: 5’-AATTCCCTGTCTTAGGTCAGTTAGGA-3’).

As an internal control we amplified samples with primers for *RPPH1*, a housekeeping gene known to have two copies per cell ^50^:

Forward: 5’-AGCTGAGTGCGTCCTGTCACT-3’

Reverse: 5’-TCTGGCCCTAGTCTCAGACCTT-3’)

The qPCR reactions were set up with 30 ng of genomic DNA (using three technical replicates), forward and reverse primers at a final concentration of 250 nM, and SYBR Green PowerUp Master Mix (ThermoFisher). Reactions were run in standard mode: a 2 min hold at 95 °C followed by 40 cycles of 15 s at 95 °C to denature and 60 s at 60 °C to anneal and extend.

### Excision efficiency determination

T cells were harvested at different time points after SB100x mRNA and viral transduction for transposase excision efficiency evaluation. Primers 5’-ccgcacgcgttctagact-3’ targeting AAV backbone and 5’-acaaagtagatgtcctaactgacttgcc-3’ targeting SB left arm were designed to evaluate SB left arm excision efficiency. Primers 5’-gccgctcggtccgcacgtg-3’ targeting AAV backbone and 5’-agtgagtttaaatgtatttggctaaggtgtatg-3’ targeting SB right arm were designed to evaluate SB right arm excision efficiency. The SYBR Green master Mix (ThermoFisher) was applied for qPCR quantification as previously described. For the excision efficiency calculation, AAV-SB-CAR only group (only transduced with AAV) was determined as baseline level of viral copy number that was existed in the T cells, then viral copy number in AAV-SB-CAR + SB100x mRNA group was divided by the baseline viral copy number, which was determined as excision efficiency.

### Flow cytometry

T cells (or other immune cells) were collected and spun down to remove media. For CAR constructs lacking a Flag tag (e.g. for CD22 and BCMA CARs), cells were incubated with CD22-Fc or BCMA-Fc protein (R&D Systems) in PBS for 30 min on ice, then stained with anti-human IgG Fc-PE and other immune markers antibodies and incubated on ice for 30 min. For CAR constructs containing a Flag tag, Flag was stained directly with an anti-Flag antibody. For the CD19.20.CAR detection, cells were incubated with biotinylated protein L (R&D) on ice for 30 min, then stained with APC streptavidin for 30 min on ice. General T cell marker staining such as exhaustion and memory followed a previous study ^78^. In brief, T cells or CAR-T cells were collected and washed with MACS buffer (0.5 % BSA and 2 mM EDTA in PBS), then stained with specific antibodies for 30 min on ice. Cells were washed with MACS buffer after staining and then analyzed on a BD FACS Aria cytometer. Data analysis was performed using FlowJo software 9.9.4 (Threestar, Ashland, OR). All flow cytometry antibodies were purchased from BioLegend. In this study, CD4+ cells refer to cell populations that were gated CD3 positive and CD8 negative. CD8+ cells refer to cell populations that were gated CD3 positive and CD8 negative.

### Kill assay (co-culture cytotoxicity)

To interrogate AAV-SB-CAR T cell killing efficacy, 1e5 of NALM6-GL (GFP-Luciferase), 1e5 of MM.1R-GL, and 5e4 of MCF7-PL (Puromycin-Luciferase) cancer cell lines per well per 100μL T cell media were seeded in 96-well plates. Corresponding CAR-T or CAR-NK cells were then added according to various effector to target (T/NK cell : cancer cell) ratios. All input CAR-T or CAR-NK cells were normalized by CAR-positive percentage to make sure each well received a consistent number of CAR-positive cells. Cytolysis was measured through luciferase assays. 150 μg / mL D-Luciferin (PerkinElmer) was added to the plate using a multi-channel pipette. Following a ∼5 min incubation at room temperature, luciferin intensity was measured by a Plate Reader (PerkinElmer).

### *In vivo* CAR-T cell functionality testing

NSG mice were intravenously injected with 5e5 NALM6-GL cancer cells. After four days of cancer inoculation, 5e6 CD22.CAR T cells were tail vein injected as treatments. Bioluminescent imaging was performed via IVIS system to monitor leukemia progression. Animal survival study followed an approved death-as-endpoint protocol.

### *In vitro* mRNA transcription

There were two sources for the mRNA used in this study: 1) commercial synthesis by TriLink Biotechnologies (TriLink mRNA was used in Fig. 1l, m; Fig. 4; Fig. S2; Fig. S3; Fig. S4 a, b; Fig. S6; Fig. S7) and 2) *in vitro* transcription (used in experiments in figures otherwise) from the SB100x plasmid using the HiScribe T7 ARCA mRNA (with tailing) Kit (NEB). Following RNA transcription *in vitro*, DNase treatment, and poly-A tailing, RNA purification was conducted using the Monarch RNA Cleanup Kit (50ug) (NEB). After the concentration of the product was measured via Nandrop (with default RNA settings), the RNA was aliquoted and stored in −80 °C. RNA was thawed on ice shortly before use in electroporation.

### Gene transfer / CAR delivery into human immune cells

The vast majority of electroporation experiments were done using a Neon system (ThermoFisher). Before electroporation, cells were collected, washed, and counted. 5e^5^-3e^6^ cells were used per reaction, depending on the specific experiment. Per 1 million cells, 1μg of SB100x mRNA was used. 100 μL Buffer R with the cell and SB100x mRNA mixture was loaded into the Neon Pipette, carefully avoiding the production of bubbles. The electroporation parameter was set at 1600 V, 10 ms, and 3 pulses for T cells, THP-1 cells, and NK-92 cells; 1900 V, 30 ms, and 1 pulse for macrophages. Cells were immediately transferred to a 24-well plate with pre-warmed media after electroporation. Depending on the experiment, specific quantities of AAV were then added to the cells at defined time points vs. electroporation (details in each figure panel and the panel’s legend, mostly 0h if not specified otherwise). Electroporation using Maxcyte system follows a similar procedure except with the manufacturer’s suggested electroporation presets. Lentiviral transduction followed standard protocols, mostly following a previous study ^78^, with the conditions specified in the figure legends.

### Insertion site library preparation and sequencing

To conduct integration site analysis for the MAJESTIC method, we created a custom protocol to prepare libraries of insertion sites from a CAR-T generation experiment: CD8+ donor 4003 T cells (collected d14 after electroporation). The custom procedure was created by combining elements of protocols from Illumina’s NEBNext® Ultra™ II FS DNA Library Prep Kit and Friedrich et al. (2017), with oligos obtained from the latter. From the non-sorted pool of T cells, roughly 10e6 CAR-Ts were collected. Then, genomic DNA was isolated using the QIAGEN Blood Mini Kit. DNA concentrations were quantified via Nanodrop. 500ng of genomics DNA were distributed into three separate tubes to serve as three technical replicates for further library preparations. Next, DNA was fragmented for 20 minutes using the Ultra II FS Enzyme Mix from NEB. 100uM Splinkerette V1.2TS and V1.2BS oligos from IDT were annealed in an Eppendorf tube by heating the mixture to 98 C for 10 min in a heat block and then unplugging the heat block to allow the reaction to cool to room temperature. The final 15uM annealed Splinkerette adaptor was ligated to the fragmented DNA reactions for 15 mins at 20 °C using NEBNext Ultra II Ligation Master Mix and Ligation Enhancer. Size selection was performed to achieve an insert size distribution of roughly 200-350bp using NEBNext Sample Purification beads: 30uL for 1^st^ bead selection 15uL for 2^nd^ selection.

Then a two-step PCR was performed, using NEBNext Ultra II Q5 Master Mix as the reaction buffer. The first PCR (98 °C 30s for one cycle; 98 °C 10s, 65 °C 75s for 18 cycles; and 65 °C 5 min for one cycle) was used to amplify genomic fragments containing the SB-left arm using two oligos: one specific to the Splinkerette adaptor, and another specific to the SB-left arm. The second PCR (98 °C 30s for one cycle; 98 °C 10s, 65 °C 75s for 12 cycles; and 65 °C 5 min for one cycle) was used to attach i7 index to the library. After each PCR, PCR cleanup using SPRIselect Purification Beads was performed. QC of each key step was performed by running 1uL of sample on a Tapestation. To quantify each library, aliquots were first diluted 1:10,000. A 1nM to 0.01pM dilution series of Illumina PhiX library was then prepared to serve as a standard. The qPCR reaction was prepared using 2x PowerUp SYBR Green, qPCR2.1 and qPCR 2.2 primers at a final concentration of 250nM, and 5uL of the diluted libraries in 20uL total volume. Quantified libraries were diluted to 2nM and then pooled in equal volumes and denatured according to the Miseq System Denature and Dilute libraries Guide. PhiX was spiked in at 50%, and the denatured pool was diluted to 8pM and sequenced on Miseq system. A 300-cycles Miseq v2 kit was used to sequence the library with a single-end setting (150 cycles for R1 and 8 cycles for the index1) Two custom sequencing primers (Spl_tag_seq for the index tag and SB_R_pr_seq for the forward read of SleepyBeauty left-end libraries) were spiked into the illumine sequencing primers according to Illumina’s bulletin on *“*Spiking custom primers into the Illumina sequencing primers” (https://support.illumina.com/bulletins/2016/04/spiking-custom-primers-into-the-illumina-sequencing-primers-.html)

### Splinkerette data processing, analysis, and visualization

Single end FASTQ reads were quality trimmed with BBDuk ^79^ using the settings trimq=27 minlen=80 maq=30 qtrim=rl. Then, non-integrated AAV-SB sequences were removed by using a sequence specific to the AAV ITR and by using Cutadapt ^80^ with the following settings: -g TATAGTCTAGAACGCGTGCG -e 0.1 --overlap 15 --discard-trimmed. Then, to trim out the transposon arms and keep only the sequences that were trimmed, we used Cutadapt with the settings -g ^ACTTCAACTG -e 0.1 -m 15 --overlap 10 --discard-untrimmed. Ten bases were removed from the 5’ end (cutadapt -u 10) and the all reads were trimmed to a fixed length of 30 (cutadapt -l 30). Reads were mapped using hisat2 ^81^ onto the HISAT2 indexed GRCh38 genome (obtained from http://daehwankimlab.github.io/hisat2/download/). Samtools view was used to filter out mapped reads with a quality score of less than 30, and the files were subsequently converted to the BED format using samtools view ^82^ and bedtools bamtobed ^83^. Genomic coordinate files in .bed format were loaded into R, keeping only the starting genomic coordinate. They were further processed and formatted into GRanges objects for data visualization. Key packages used for R processing and visualization include GenomicRanges^84^, genomation^85^, ggbio^86^, BRGenomics (https://mdeber.github.io), and pheatmap (https://cran.r-project.org/web/packages/pheatmap/pheatmap.pdf).

### Sample size determination

Sample size was determined according to the lab’s prior work, cited literature, or similar approaches in the field.

### Replication

Experimental replications are indicated in each figure panel’s legend. Important experiments have been repeated independently to ensure reproducibility.

### Randomization and blinding statements

*In vitro* cell culture experiments were not randomized. Investigators were not blinded in *in vitro* cell culture experiments. Mouse experiments were randomized by using littermates, and blinded using generic cage barcodes and eartags where applicable.

### Data Collection summary

Flow cytometry data was collected by BD FACSAria.

Co-culture killing assay data were collected with Perkin Elmer Envision Plate Reader.

### Data analysis summary

Flow cytometry data were analyzed by FlowJo v.10.7.

All simple statistical analyses were done with Prism 9.

All NGS analyses were performed using custom bash and R scripts.

### Standard statistical analysis

Various standard non-NGS statistical analyses were performed. All statistical methods are described in figure legends and/or supplementary Excel tables. The *p* values and statistical significance were estimated for data was sourced from different donors. Different levels of statistical significance were accessed based on specific *p* values and type I error cutoffs (0.05, 0.01, 0.001, 0.0001). Standard analysis was performed using GraphPad Prism. NGS statistics were performed using custom bash and R scripts.

### Code availability

Scripts used to process the insertion site mapping data will be available at https://github.com/stanleyzlam/SB-CAR.

### Data and resource availability

Data generated or analyzed during this study are included in this published article (and its supplementary information files). Specifically, source data and statistics are provided in an Excel file. Raw and processed genome sequence data from the Splinkerette experiments will be deposited to public databases such as NIH Sequence Read Archive (SRA) / Gene Expression Omnibus (GEO) under accession number (GSE220202, token mtqrayyonvwfzmd). Data and materials that support the findings of this research are available to the academic community from the corresponding author upon reasonable request, via direct sharing, MTAs, or public repositories.

## Figure notes

Data are shown as mean ± s.e.m., with individual data points in plots, unless otherwise noted. Significance notes: ns - not significant; * p < 0.05, ** p < 0.01, *** p < 0.001, **** p < 0.0001. Source data, statistical methods and additional statistics for non-high-throughput experiments are provided in a supplemental Excel file.

## Supplemental Materials

### Supplemental Figure Legends

**Figure S1.**
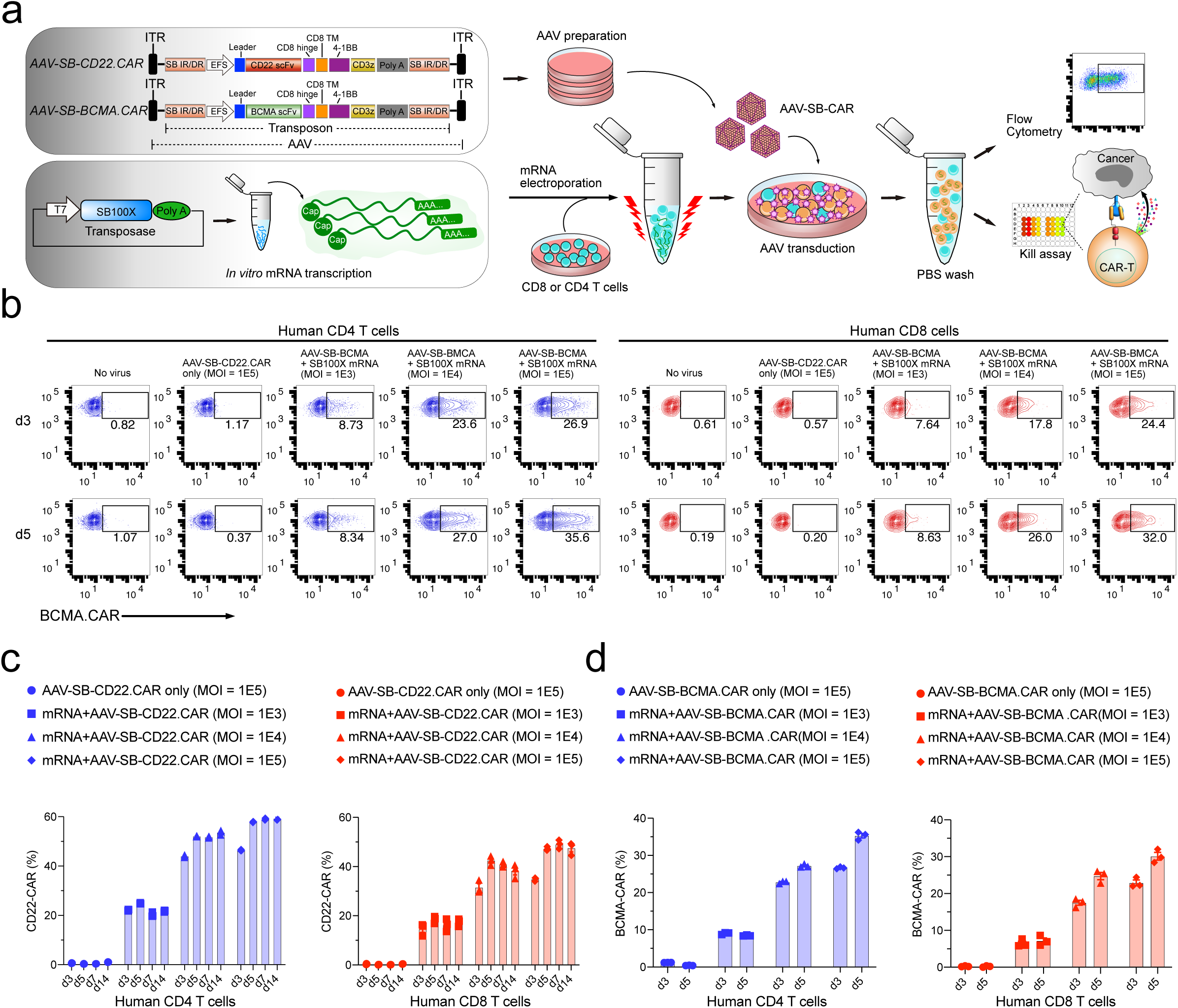
MAJESTIC CAR generation and optimization. **(a)** Schematic of AAV-SB-CD22.CAR, AAV-SB-BCMA.CAR, and SB100X constructs and key procedures: mRNA *in vitro* transcription, AAV production, mRNA electroporation, flow cytometry, and kill assay. **(b)** Representative flow cytometry plots of AAV-SB-BCMA.CAR CD4 (left) and CD8 (right) T cells show the percentage of CAR-expressing cells. CD4 cells are defined as CD3^+^ and CD8^-^ cells. **(c)** Quantification of CD22.CAR T cell ratio of (fig.1b, c). **(d)** Quantification of BCMA.CAR T ratio in human CD4 and CD8 T cells. In this figure, for optimization of conditions, each assay was done with one donor with three technical replicates. Donor 2 T cells were used in this figure.

**Figure S2.**
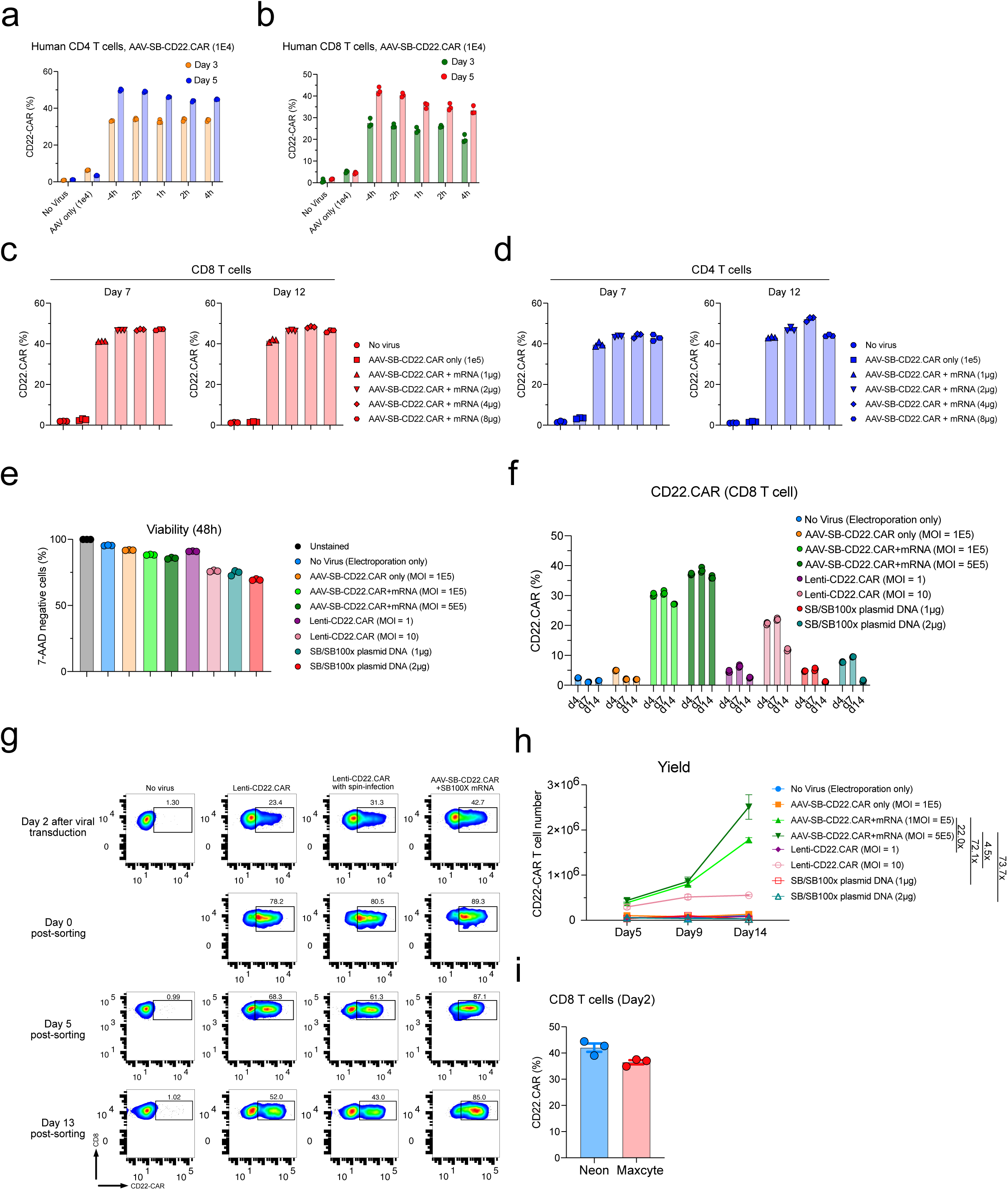
Time point optimization, SB100X transposase mRNA titration, and MAJESTIC CAR T yield. **(a)** Quantification of CD22.CAR T cell ratio for CD4 T cells that were transduced with AAV-SB-CD22.CAR virus at various time points. **(b)** Quantification of CD22.CAR T cell ratio for CD8 T cells that were transduced with AAV-SB-CD22.CAR virus at various time points. **(c, d)** Representative flow cytometry plots of CD22.CAR T cells produced via AAV-SB and a titrated serial of SB100X mRNA. **(c)** CD8 T cells. (**d**) CD4 T cells. **(e)** Quantification of the cell viability of CD8 T cells. **(f)** Quantification of CAR T cells. **(g)** Flow cytometry plots of CAR T cell ratios before and after sorting. **(h)** CD22.CAR T cell yield quantification (yield = total viable cell count x CAR-positive percentage). Cells were split into 3 technical replicates after electroporation. Yield is calculated for each technical replicate separately. **(i)** CAR+ T cell generation efficiency (CAR+%) of MAJESTIC using Neon and Maxcyte platforms. In this figure, for optimization of conditions, each assay was done with one donor with three technical replicates. Donor 2 and donor 0286 T cells were used in this figure.

**Figure S3.**
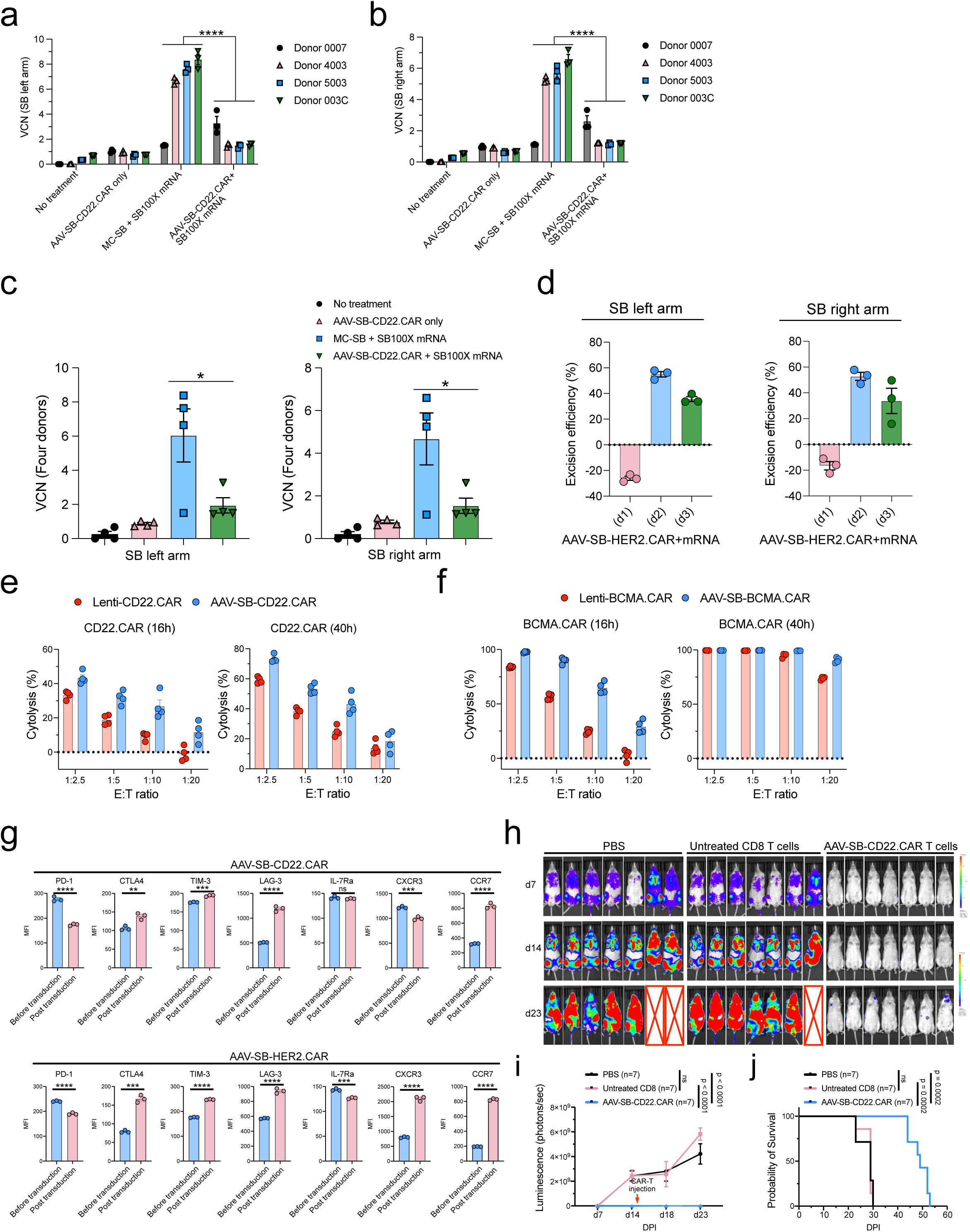
Vector copy number quantification, immune marker profiling, and functionality testing of MAJESTIC-produced CD8 CAR-T cells. **(a-c)** Vector copy number (VCN) quantification of MAJESTIC-manufactured CAR-T cells. Purified CAR T cells were collected for DNA extraction after three weeks of mRNA electroporation and viral transduction. (a) left arm probe, (b) right arm probe, (c) left panel: left arm probe, right panel: right arm probe. **(d)** SB100X transposase excision efficiency evaluation. Left panel: left arm probe, right panel: right arm probe. **(e)** Cytolysis analysis of NAML6-GL (NAML6 with GFP and luciferase reporters) cancer cells that were co-cultured with Lenti-CD22.CAR and AAV-SB-CD22.CAR T cells. CAR-Ts were seeded at various effector : target (E:T) ratios, and luciferase imaging was performed at two time points (16h and 40h). **(f)** Cytolysis analysis of MM.1R-GL (MM.1R with GFP and luciferase reporters) cancer cells that were co-cultured with Lenti-BCMA.CAR and AAV-SB-BCMA.CAR T cells. CAR-Ts were seeded at various effector : target (E:T) ratios, and luciferase imaging was performed at two time points (16h and 40h). **(g)** Exhaustion and memory marker expression in CD22-CAR and HER2-CAR T cells before and post transfection. Unpaired *t* tests were performed to evaluate statistical significance. **(h)** Bioluminescent density of *NSG* mice that were injected with NALM6-GL cancer cells and with CD22-CAR therapy (n = 7 mice per group). **(i)** Quantification of total luminescence for (h). n = 7 mice. Two-way ANOVA with multiple comparisons tests was performed to evaluate statistical significance. **(j)** Survival curve of NALM6-GL-induced leukemia-bearing NSG mice that treated with PBS, untreated CD8 T cells, and AAV-SB-CD22.CAR T cells. Log-rank (Mantel-Cox) tests were performed to evaluate statistical significance. Donor 0007, 4003, 5003, 003C, 0286 T cells were used in this figure.

**Figure S4.**
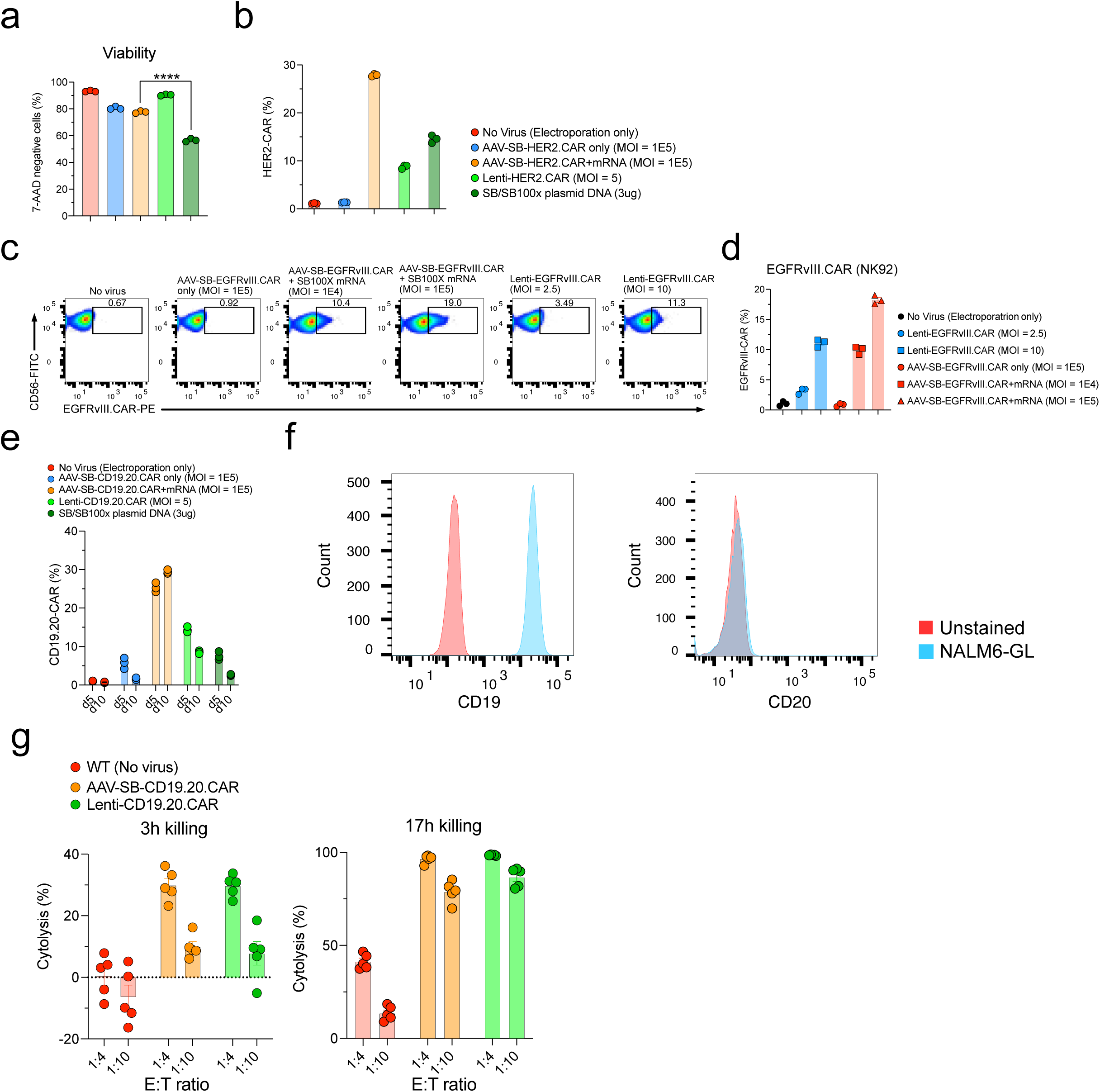
Generation of various therapeutic immune cells by MAJESTIC. **(a)** Quantification of the HER2.CAR T cell viability. **(b)** Quantification of HER2.CAR-positive CD8 T cells. **(c)** Representative flow cytometry plots of EGFRvIII.CAR-NK92 cells were produced via either the SB system or lentiviral transduction. **(d)** Quantification of (c). **(e)** Quantification of CD19.20.CAR T cells. **(f)** Flow cytometry detection of CD19 and CD20 expression in NALM6-GL cells. **(g)** Cytolysis analysis of NALM6-GL cancer cells that were co-cultured with lenti-CD19.20.CAR and AAV-SB-CD19.20.CAR T cells. Donor 2 and donor VP2 T cells were used in this figure.

**Figure S5.**
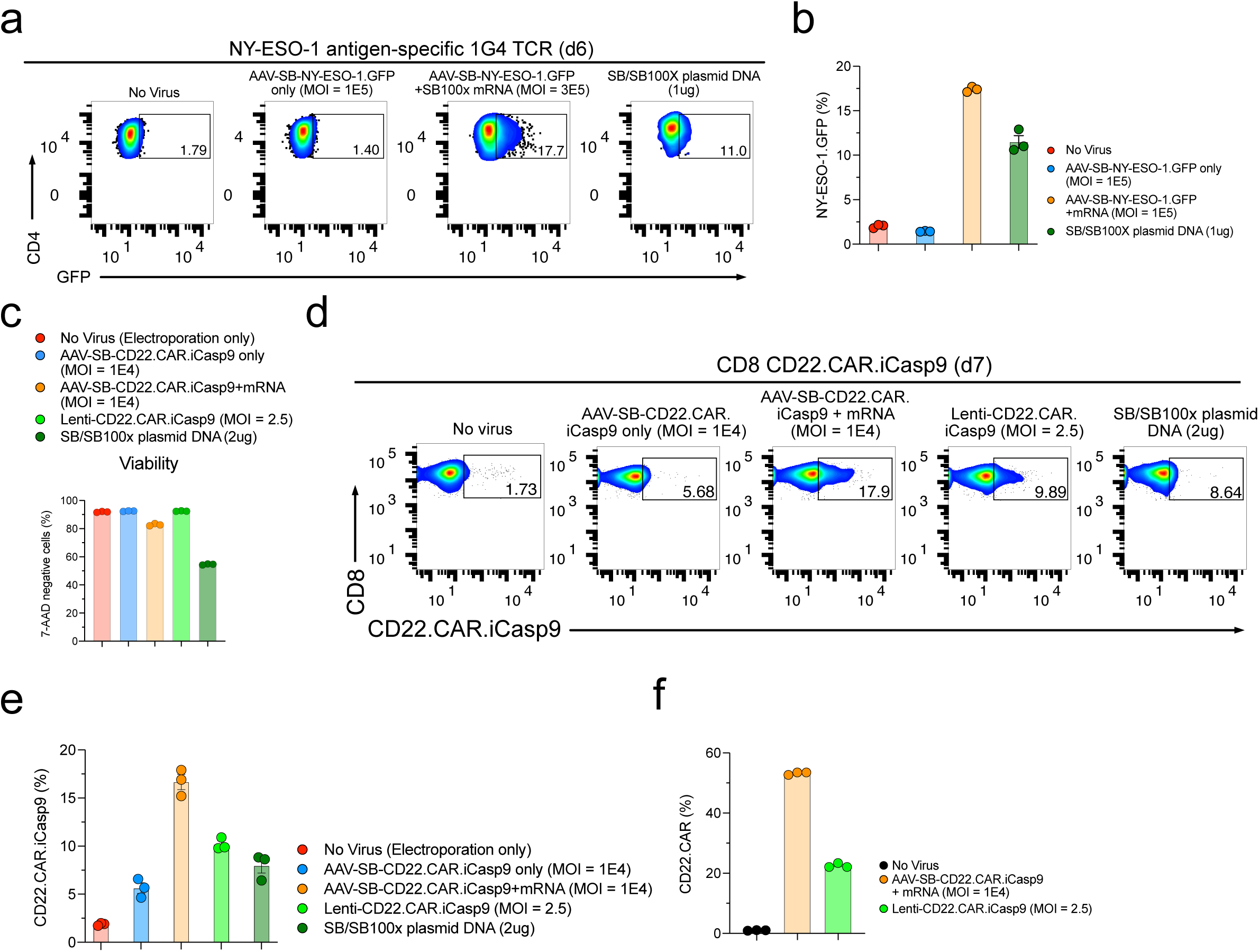
Generation of various therapeutic immune cells by MAJESTIC. **(a)** Representative flow cytometry plots of NY-ESO-1 T cells. **(b)** Quantification of (a). **(c)** Quantification of the CD22.CAR.iCasp T cell viability. **(d)** Representative flow cytometry plots of CD22.CAR.iCasp9 T cells. **(e)** Quantification of (d). **(f)** Quantification of the CD22.CAR.iCasp9 T cells post antigen-specific cancer cells stimulation. In this figure, each assay has three technical replicates, donor 601c and donor 02 T cells were used in this figure.

**Figure S6.**
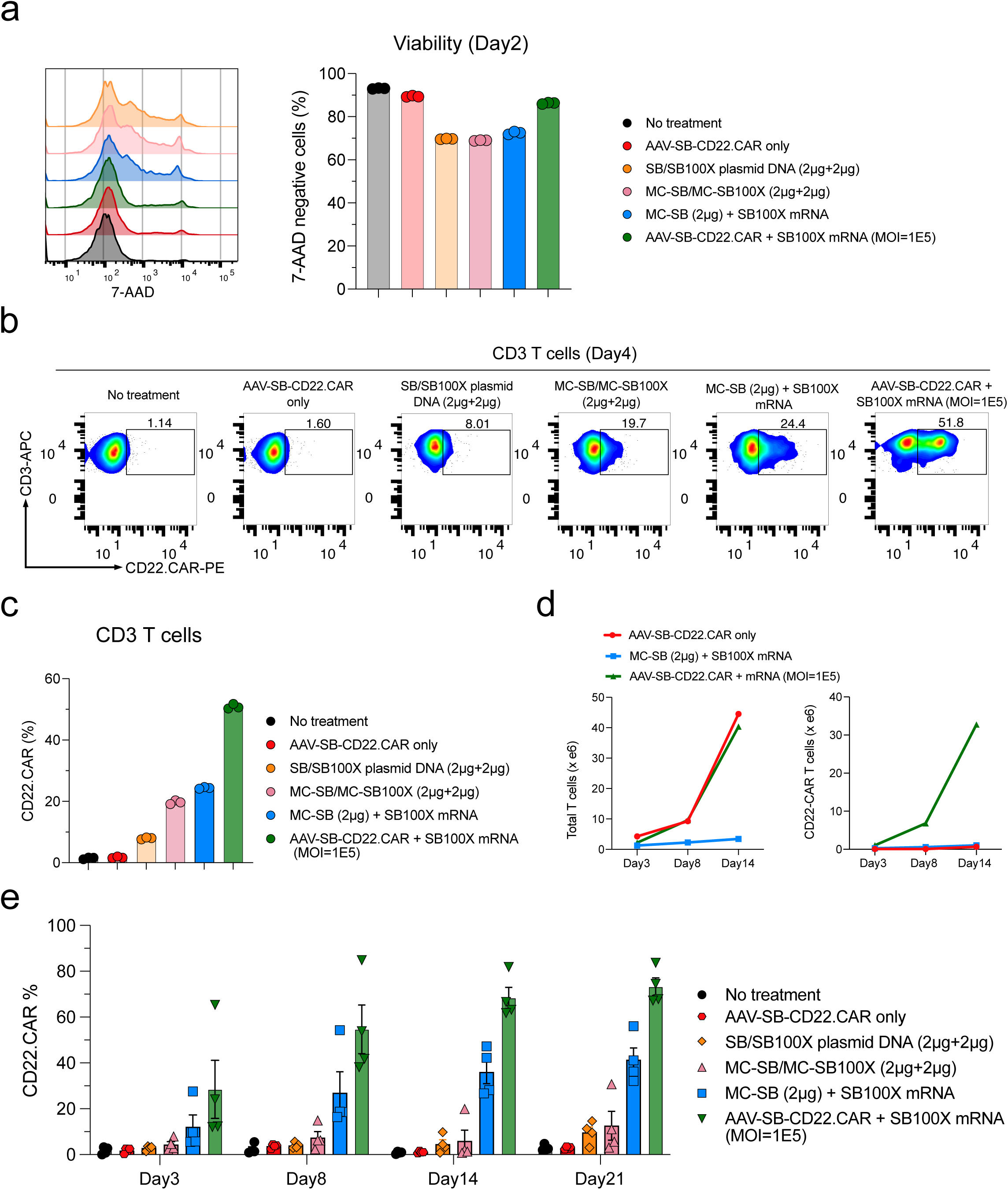
Gene delivery efficiency comparison of MC-SB/SB100X mRNA with the MAJESTIC system. **(a)** Flow cytometry histogram overlays and bar plots of cell viability post-electroporation as measured with 7-AAD staining. **(b)** Flow cytometry data of CD22.CAR T cells from human primary CD3 T cells produced by plasmid transposon plasmid, transposon MC, transposon MC with mRNA-transposase, and MAJESTIC. **(c)** Quantification of (b). **(d)** Yield calculations (yield = CAR% * total viable cell count). All conditions started with an equal amount of primary T cells per replicate. Three CAR% replicates were averaged and then multiplied by the average of 2 cell count replicates. Left panel, total T cell count; Right panel, total CAR+ T cell count. **(e)** Quantification of flow cytometry data of CD22.CAR T cells from four human PBMCs (same data as Fig. 4 e-f, plotted in dot-whisker plots).

**Figure S7.**
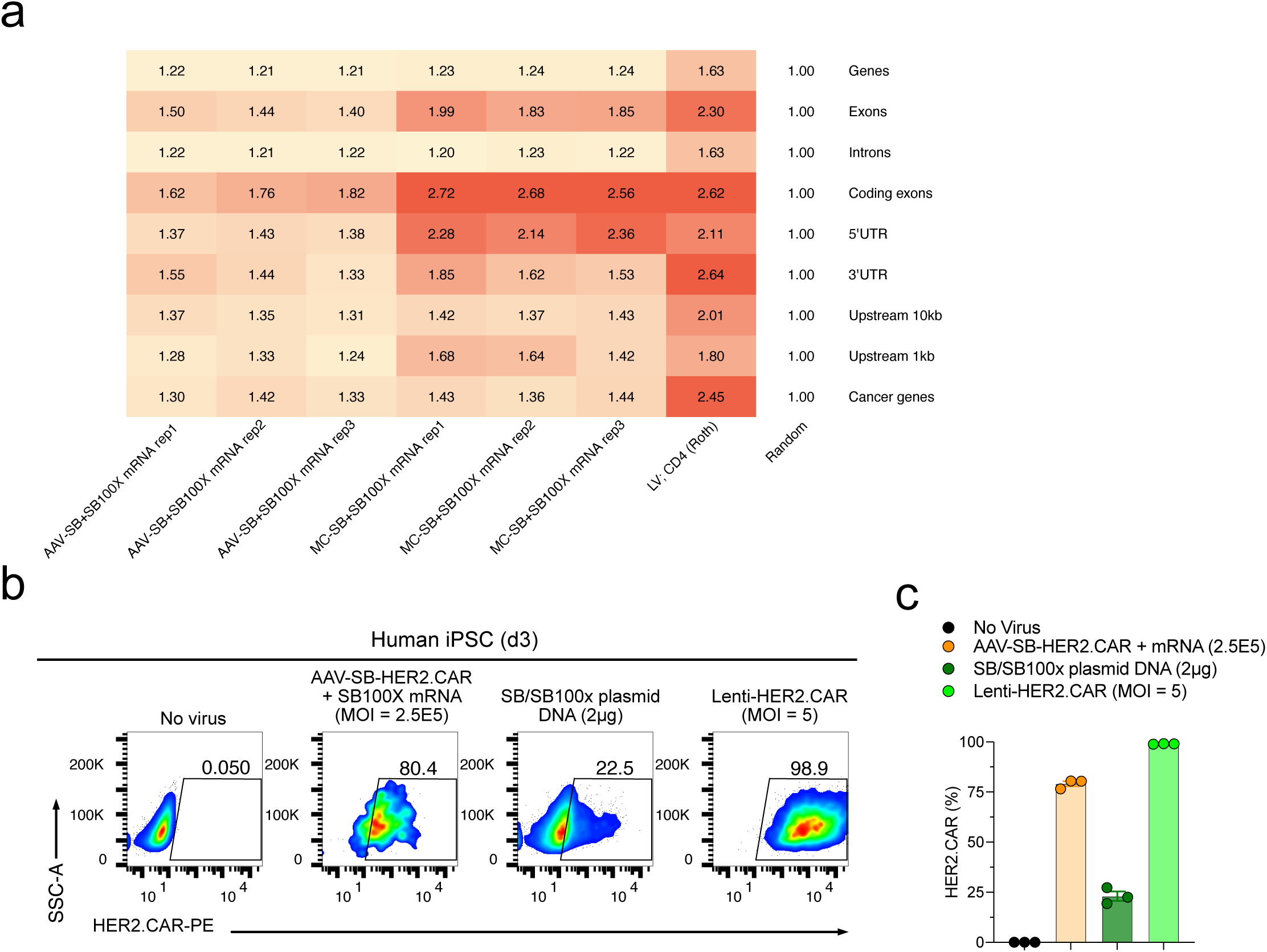
Genomic integration of the *Sleeping Beauty* transposon and iPSC-CAR generation. **(a)** A heatmap of relative quantifications of insertions in functional gene regions for various gene transfer methods. For each condition, the proportion of reads falling into a given gene region is first calculated. Then a fold-change relative to the frequency of randomly generated insertions into that same region produces the value shown in the heatmap. UTR = untranslated regions. The list of cancer genes was taken from COSMIC and included genes from both Tier 1 and Tier 2 (https://cancer.sanger.ac.uk/census). **(b)** Representative flow cytometry plots of iPSC-HER2.CAR cells. **(c)** Quantification of (b).

### Source Data and Statistics

Non-NGS source data and statistics for all data in the figures.

### Supplemental Tables

Table S1. List of key oligos used

## References

1. Hayes, C. Cellular immunotherapies for cancer. Ir J Med Sci 190, 41–57 (2021).

2. Laskowski, T. & Rezvani, K. Adoptive cell therapy: Living drugs against cancer. J Exp Med 217 (2020).

3. June, C.H., O’Connor, R.S., Kawalekar, O.U., Ghassemi, S. & Milone, M.C. CAR T cell immunotherapy for human cancer. Science 359, 1361–1365 (2018).

4. Majzner, R.G. & Mackall, C.L. Clinical lessons learned from the first leg of the CAR T cell journey. Nat Med 25, 1341–1355 (2019).

5. Upadhaya, S. et al. The clinical pipeline for cancer cell therapies. Nat Rev Drug Discov 20, 503–504 (2021).

6. Verdegaal, E.M. et al. Neoantigen landscape dynamics during human melanoma-T cell interactions. Nature 536, 91–95 (2016).

7. Johnson, L.A. et al. Gene therapy with human and mouse T-cell receptors mediates cancer regression and targets normal tissues expressing cognate antigen. Blood 114, 535–546 (2009).

8. Thomas, R. et al. NY-ESO-1 Based Immunotherapy of Cancer: Current Perspectives. Front Immunol 9, 947 (2018).

9. Chu, J. et al. CS1-specific chimeric antigen receptor (CAR)-engineered natural killer cells enhance in vitro and in vivo antitumor activity against human multiple myeloma. Leukemia 28, 917–927 (2014).

10. Genssler, S. et al. Dual targeting of glioblastoma with chimeric antigen receptor-engineered natural killer cells overcomes heterogeneity of target antigen expression and enhances antitumor activity and survival. Oncoimmunology 5, e1119354 (2016).

11. Zhang, Q. et al. Synergistic Effects of Cabozantinib and EGFR-Specific CAR-NK-92 Cells in Renal Cell Carcinoma. J Immunol Res 2017, 6915912 (2017).

12. Kruschinski, A. et al. Engineering antigen-specific primary human NK cells against HER-2 positive carcinomas. Proc Natl Acad Sci U S A 105, 17481–17486 (2008).

13. Liu, E. et al. Cord blood NK cells engineered to express IL-15 and a CD19-targeted CAR show long-term persistence and potent antitumor activity. Leukemia 32, 520–531 (2018).

14. Chu, Y. et al. Targeting CD20+ Aggressive B-cell Non-Hodgkin Lymphoma by Anti-CD20 CAR mRNA-Modified Expanded Natural Killer Cells In Vitro and in NSG Mice. Cancer Immunol Res 3, 333–344 (2015).

15. Li, Y., Hermanson, D.L., Moriarity, B.S. & Kaufman, D.S. Human iPSC-Derived Natural Killer Cells Engineered with Chimeric Antigen Receptors Enhance Anti-tumor Activity. Cell Stem Cell 23, 181–192 e185 (2018).

16. Klichinsky, M. et al. Human chimeric antigen receptor macrophages for cancer immunotherapy. Nat Biotechnol 38, 947–953 (2020).

17. Zhang, L. et al. Pluripotent stem cell-derived CAR-macrophage cells with antigen-dependent anti-cancer cell functions. J Hematol Oncol 13, 153 (2020).

18. Themeli, M. et al. Generation of tumor-targeted human T lymphocytes from induced pluripotent stem cells for cancer therapy. Nat Biotechnol 31, 928–933 (2013).

19. Di Stasi, A. et al. Inducible apoptosis as a safety switch for adoptive cell therapy. N Engl J Med 365, 1673–1683 (2011).

20. Shah, N.N. et al. Bispecific anti-CD20, anti-CD19 CAR T cells for relapsed B cell malignancies: a phase 1 dose escalation and expansion trial. Nat Med 26, 1569–1575 (2020).

21. Morgan, R.A. & Boyerinas, B. Genetic Modification of T Cells. Biomedicines 4 (2016).

22. Ellis, J. Silencing and variegation of gammaretrovirus and lentivirus vectors. Hum Gene Ther 16, 1241–1246 (2005).

23. Ciuffi, A. Mechanisms governing lentivirus integration site selection. Curr Gene Ther 8, 419–429 (2008).

24. Hacein-Bey-Abina, S. et al. LMO2-associated clonal T cell proliferation in two patients after gene therapy for SCID-X1. Science 302, 415–419 (2003).

25. Hacein-Bey-Abina, S. et al. Insertional oncogenesis in 4 patients after retrovirus-mediated gene therapy of SCID-X1. J Clin Invest 118, 3132–3142 (2008).

26. Kustikova, O. et al. Clonal dominance of hematopoietic stem cells triggered by retroviral gene marking. Science 308, 1171–1174 (2005).

27. Zhou, S. et al. Evaluating the Safety of Retroviral Vectors Based on Insertional Oncogene Activation and Blocked Differentiation in Cultured Thymocytes. Mol Ther 24, 1090–1099 (2016).

28. Naso, M.F., Tomkowicz, B., Perry, W.L., 3rd & Strohl, W.R. Adeno-Associated Virus (AAV) as a Vector for Gene Therapy. BioDrugs 31, 317–334 (2017).

29. Eyquem, J. et al. Targeting a CAR to the TRAC locus with CRISPR/Cas9 enhances tumour rejection. Nature 543, 113–117 (2017).

30. Dai, X. et al. One-step generation of modular CAR-T cells with AAV-Cpf1. Nat Methods 16, 247–254 (2019).

31. Nahmad, A.D. et al. Frequent aneuploidy in primary human T cells after CRISPR-Cas9 cleavage. Nat Biotechnol (2022).

32. Papathanasiou, S. et al. Whole chromosome loss and genomic instability in mouse embryos after CRISPR-Cas9 genome editing. Nat Commun 12, 5855 (2021).

33. Zuccaro, M.V. et al. Allele-Specific Chromosome Removal after Cas9 Cleavage in Human Embryos. Cell 183, 1650–1664 e1615 (2020).

34. Wilson, M.H., Coates, C.J. & George, A.L., Jr. PiggyBac transposon-mediated gene transfer in human cells. Mol Ther 15, 139–145 (2007).

35. Doherty, J.E. et al. Hyperactive piggyBac gene transfer in human cells and in vivo. Hum Gene Ther 23, 311–320 (2012).

36. Kebriaei, P., Izsvak, Z., Narayanavari, S.A., Singh, H. & Ivics, Z. Gene Therapy with the Sleeping Beauty Transposon System. Trends Genet 33, 852–870 (2017).

37. Sebastian-Martin, A., Barrioluengo, V. & Menendez-Arias, L. Transcriptional inaccuracy threshold attenuates differences in RNA-dependent DNA synthesis fidelity between retroviral reverse transcriptases. Sci Rep 8, 627 (2018).

38. Monjezi, R. et al. Enhanced CAR T-cell engineering using non-viral Sleeping Beauty transposition from minicircle vectors. Leukemia 31, 186–194 (2017).

39. Singh, H. et al. Manufacture of clinical-grade CD19-specific T cells stably expressing chimeric antigen receptor using Sleeping Beauty system and artificial antigen presenting cells. PLoS One 8, e64138 (2013).

40. Magnani, C.F. et al. Sleeping Beauty-engineered CAR T cells achieve antileukemic activity without severe toxicities. J Clin Invest 130, 6021–6033 (2020).

41. Chicaybam, L. et al. CAR T Cells Generated Using Sleeping Beauty Transposon Vectors and Expanded with an EBV-Transformed Lymphoblastoid Cell Line Display Antitumor Activity In Vitro and In Vivo. Hum Gene Ther 30, 511–522 (2019).

42. Geurts, A.M. et al. Gene mutations and genomic rearrangements in the mouse as a result of transposon mobilization from chromosomal concatemers. PLoS Genet 2, e156 (2006).

43. Liu, M.A. A Comparison of Plasmid DNA and mRNA as Vaccine Technologies. Vaccines (Basel) 7 (2019).

44. Ye, L. et al. In vivo CRISPR screening in CD8 T cells with AAV-Sleeping Beauty hybrid vectors identifies membrane targets for improving immunotherapy for glioblastoma. Nat Biotechnol 37, 1302–1313 (2019).

45. Mates, L. et al. Molecular evolution of a novel hyperactive Sleeping Beauty transposase enables robust stable gene transfer in vertebrates. Nat Genet 41, 753–761 (2009).

46. Francois, A. et al. Accurate Titration of Infectious AAV Particles Requires Measurement of Biologically Active Vector Genomes and Suitable Controls. Mol Ther Methods Clin Dev 10, 223–236 (2018).

47. Ghorashian, S. et al. Enhanced CAR T cell expansion and prolonged persistence in pediatric patients with ALL treated with a low-affinity CD19 CAR. Nat Med 25, 1408–1414 (2019).

48. Chicaybam, L. et al. Transposon-mediated generation of CAR-T cells shows efficient anti B-cell leukemia response after ex vivo expansion. Gene Ther 27, 85–95 (2020).

49. Xia, X., Zhang, Y., Zieth, C.R. & Zhang, S.C. Transgenes delivered by lentiviral vector are suppressed in human embryonic stem cells in a promoter-dependent manner. Stem Cells Dev 16, 167–176 (2007).

50. Kolacsek, O. et al. Reliable transgene-independent method for determining Sleeping Beauty transposon copy numbers. Mob DNA 2, 5 (2011).

51. Schneider, D. et al. A tandem CD19/CD20 CAR lentiviral vector drives on-target and off-target antigen modulation in leukemia cell lines. J Immunother Cancer 5, 42 (2017).

52. Straathof, K.C. et al. An inducible caspase 9 safety switch for T-cell therapy. Blood 105, 4247–4254 (2005).

53. Sadelain, M., Papapetrou, E.P. & Bushman, F.D. Safe harbours for the integration of new DNA in the human genome. Nat Rev Cancer 12, 51–58 (2011).

54. Roth, S.L., Malani, N. & Bushman, F.D. Gammaretroviral integration into nucleosomal target DNA in vivo. J Virol 85, 7393–7401 (2011).

55. Hou, A.J., Chen, L.C. & Chen, Y.Y. Navigating CAR-T cells through the solid-tumour microenvironment. Nat Rev Drug Discov 20, 531–550 (2021).

56. Dolgin, E. Cancer-eating immune cells kitted out with CARs. Nat Biotechnol 38, 509–511 (2020).

57. Marofi, F. et al. CAR-NK Cell: A New Paradigm in Tumor Immunotherapy. Front Oncol 11, 673276 (2021).

58. Bailey, S.R. & Maus, M.V. Gene editing for immune cell therapies. Nat Biotechnol 37, 1425–1434 (2019).

59. Tang, X. et al. First-in-man clinical trial of CAR NK-92 cells: safety test of CD33-CAR NK-92 cells in patients with relapsed and refractory acute myeloid leukemia. Am J Cancer Res 8, 1083–1089 (2018).

60. Auwerx, J. The human leukemia cell line, THP-1: a multifacetted model for the study of monocyte-macrophage differentiation. Experientia 47, 22–31 (1991).

61. Miskey, C. et al. Engineered Sleeping Beauty transposase redirects transposon integration away from genes. Nucleic Acids Res 50, 2807–2825 (2022).

62. Querques, I. et al. A highly soluble Sleeping Beauty transposase improves control of gene insertion. Nat Biotechnol 37, 1502–1512 (2019).

63. Roth, T.L. et al. Reprogramming human T cell function and specificity with non-viral genome targeting. Nature 559, 405–409 (2018).

64. Fu, Y. et al. High-frequency off-target mutagenesis induced by CRISPR-Cas nucleases in human cells. Nat Biotechnol 31, 822–826 (2013).

65. Anderson, K.R. et al. CRISPR off-target analysis in genetically engineered rats and mice. Nat Methods 15, 512–514 (2018).

66. Frock, R.L. et al. Genome-wide detection of DNA double-stranded breaks induced by engineered nucleases. Nat Biotechnol 33, 179–186 (2015).

67. Tsai, S.Q. et al. GUIDE-seq enables genome-wide profiling of off-target cleavage by CRISPR-Cas nucleases. Nat Biotechnol 33, 187–197 (2015).

68. Cameron, P. et al. Mapping the genomic landscape of CRISPR-Cas9 cleavage. Nat Methods 14, 600–606 (2017).

69. Tsai, S.Q. et al. CIRCLE-seq: a highly sensitive in vitro screen for genome-wide CRISPR-Cas9 nuclease off-targets. Nat Methods 14, 607–614 (2017).

70. Takata, M. et al. Homologous recombination and non-homologous end-joining pathways of DNA double-strand break repair have overlapping roles in the maintenance of chromosomal integrity in vertebrate cells. Embo J 17, 5497–5508 (1998).

71. Enache, O.M. et al. Cas9 activates the p53 pathway and selects for p53-inactivating mutations. Nat Genet 52, 662–668 (2020).

72. Zhang, W. et al. Hybrid adeno-associated viral vectors utilizing transposase-mediated somatic integration for stable transgene expression in human cells. PLoS One 8, e76771 (2013).

73. Clauss, J. et al. Efficient Non-Viral T-Cell Engineering by Sleeping Beauty Minicircles Diminishing DNA Toxicity and miRNAs Silencing the Endogenous T-Cell Receptors. Hum Gene Ther 29, 569–584 (2018).

74. Jin, Z. et al. The hyperactive Sleeping Beauty transposase SB100X improves the genetic modification of T cells to express a chimeric antigen receptor. Gene Ther 18, 849–856 (2011).

75. Kovac, A. et al. RNA-guided retargeting of Sleeping Beauty transposition in human cells. Elife 9 (2020).

76. Muther, N., Noske, N. & Ehrhardt, A. Viral hybrid vectors for somatic integration - are they the better solution? Viruses 1, 1295–1324 (2009).

77. Balciunas, D. et al. Harnessing a high cargo-capacity transposon for genetic applications in vertebrates. PLoS Genet 2, e169 (2006).

78. Ye, L. et al. A genome-scale gain-of-function CRISPR screen in CD8 T cells identifies proline metabolism as a means to enhance CAR-T therapy. Cell Metab 34, 595–614 e514 (2022).

79. 79. Bushnell, B. in 9th Annual Genomics of Energy & Environment Meeting (Walnut Creek; 2014).

80. Martin, M. Cutadapt Removes Adapter Sequences From High-Throughput Sequencing Reads. EMBnet.joural 17 (2011).

81. Kim, D., Paggi, J.M., Park, C., Bennett, C. & Salzberg, S.L. Graph-based genome alignment and genotyping with HISAT2 and HISAT-genotype. Nat Biotechnol 37, 907–915 (2019).

82. Li, H. et al. The Sequence Alignment/Map format and SAMtools. Bioinformatics 25, 2078–2079 (2009).

83. Quinlan, A.R. & Hall, I.M. BEDTools: a flexible suite of utilities for comparing genomic features. Bioinformatics 26, 841–842 (2010).

84. Lawrence, M. et al. Software for computing and annotating genomic ranges. PLoS Comput Biol 9, e1003118 (2013).

85. Akalin, A., Franke, V., Vlahovicek, K., Mason, C.E. & Schubeler, D. Genomation: a toolkit to summarize, annotate and visualize genomic intervals. Bioinformatics 31, 1127–1129 (2015).

86. Yin, T., Cook, D. & Lawrence, M. ggbio: an R package for extending the grammar of graphics for genomic data. Genome Biol 13, R77 (2012).

